# A Crystallographic Snapshot of SARS-CoV-2 Main Protease Maturation Process

**DOI:** 10.1101/2020.12.23.424149

**Authors:** G. D. Noske, A. M. Nakamura, V. O. Gawriljuk, R. S. Fernandes, G. M. A. Lima, H. V. D. Rosa, H. D. Pereira, A. C. M. Zeri, A.A. F. Z. Nascimento, M. C. L. C. Freire, G. Oliva, A. S. Godoy

## Abstract

SARS-CoV-2 is the causative agent of COVID-19. The dimeric form of the viral main protease is responsible for the cleavage of the viral polyprotein in 11 sites, including its own N and C-terminus. Although several mechanisms of self-cleavage had been proposed for SARS-CoV, the lack of structural information for each step is a setback to the understanding of this process. Herein, we used X-ray crystallography to characterize an immature form of the main protease, which revealed major conformational changes in the positioning of domain-three over the active site, hampering the dimerization and diminishing its activity. We propose that this form preludes the cis-cleavage of N-terminal residues within the dimer, leading to the mature active site. Using fragment screening, we probe new cavities in this form which can be used to guide therapeutic development. Furthermore, we characterized a serine site-directed mutant of the main protease bound to its endogenous N and C-terminal residues during the formation of the tetramer. This quaternary form is also present in solution, suggesting a transitional state during the C-terminal trans-cleavage. This data sheds light in the structural modifications of the SARS-CoV-2 main protease during maturation, which can guide the development of new inhibitors targeting its intermediary states.

## Main Text

Severe acute respiratory syndrome coronavirus 2 (SARS-CoV-2) is the causative agent of COVID-19, a highly infectious disease that rapidly spread causing a global pandemic. SARS-CoV-2 is an enveloped RNA virus belonging to the β-lineage of coronaviruses, which includes SARS-CoV and Middle East (MERS-CoV) respiratory viruses ^1–3^. The viral genome is a single-stranded positive RNA comprising about 30,000 nucleotides, that shares 82% sequence similarity with SARS-CoV ^4^. The replicase gene (ORF1ab) encodes two overlapping polyproteins (pp1a and pp1ab) that are required for viral replication and transcription ^5^. The main protease (M^pro^), also known as 3C-like protease (3CL^pro^) is a viral cysteine protease specific for glutamine at the S1 subsite, showing variable recognition preferences at S2 (Leu/Phe/Met/Val) and S2’ subsites (Ser/Ala/Gly/Asn) ^6^. M^pro^ is responsible for the maturation of pp1a and pp1ab in at least 11 characterized sites, including its auto-processing at the N and C terminus, which is essential for its activity and dimerization ^7–9^. Due to its essential role in viral replication, M^pro^ is one of the most well characterized non-structural proteins of SARS-CoV-2. In addition, its unique features of cleavage site recognition and the absence of closely related homologues in humans, identify M^pro^ as a major target for antiviral drug development ^9–11^.

Although M^pro^ activity is crucial to viral biology, its self-maturation process is still poorly understood. Several biochemical and crystallographic studies on native and mutated forms of SARS-CoV M^pro^ tried to elucidate its maturation mechanism (reviewed in ^12^), by evaluating if the N and C-terminus processing occurs within a dimer (*cis*-cleavage) or between two distinct dimers (*trans*-cleavage). The first 2005 model suggested that M^pro^ probably forms a small amount of active dimer after autocleavage that immediately enables the catalytic site to act on other cleavage sites in the polyprotein ^13^. In 2010, based on the observation that dimerization of mature M^pro^ is enhanced by the presence of substrates, Li and colleagues proposed that after the translation, two M^pro^ protomers form a transient dimer which is stabilized by binding the N-terminal site of its substrate (another M^pro^ in polyprotein) and further cleave to free its N-terminus ^14^. In addition, Chen et al. (2010) suggested that the N-terminal autocleavage might only need two immature forms of M^pro^ in monomeric polyproteins to form an intermediate dimer that is not related to the active dimer of the mature enzyme ^15^.

Herein, we used X-ray crystallography integrated with biochemical techniques to investigate the self-maturation process of SARS-CoV-2 M^pro^. The construct of M^pro^ containing N-terminal insertions produced an immature form of the enzyme (IMT M^pro^), unable to form a dimer, that showed a reduced enzymatic activity. We used fragment screening to probe new cavities for drug development in this construct. The inactive mutant C145S with inserted native N-terminal residues (C145S M^pro^) produced a form of the protein that behaves as monomers, dimers, trimers and tetramers in solution. Crystals of the tetrameric form revealed details of how M^pro^ self-processes its N and C-terminal residues. All forms of the enzyme revealed important conformation changes that can guide direct-acting drug development.

### Activity and biochemical characterization

A general strategy to produce SARS-CoV-2 M^pro^ is to maintain its self-cleavage N-terminal portion and add the HRV-3C cleavage site with a histidine-tag at the C-terminal portion. We successfully used ammonium sulfate precipitation followed by ion exchange chromatography to obtain pure M^pro^, simplifying the protocol to one that takes less than 8 h and with a final yield of ∼2.5 mg/L of culture. The SARS-CoV-2 IMT M^pro^ was obtained by adding a non-cleavable sequence (Gly-Ala-Met) at the N-terminal Ser1 of M^pro^, and purified by a similar protocol. The SARS-CoV-2 IMT M^pro^ was produced as a soluble protein, yielding ∼80 mg/L of culture. To further investigate the role of N-terminal residues in the maturation of M^pro^, we designed a construct containing the mutated C145S residue with its native cleavage peptide of M^pro^ (Ser-4,Ala-3,Val-2,Leu-1,Gln0↓) at the N-terminal of Ser1 (Fig. S1). During gel filtration, two M^pro^ peaks were identified with mass consistent with a monomer and a tetramer (Fig. S1).

M^pro^ and IMT M^pro^ demonstrate to be active and able to recognize and cleave the fluorogenic substrate (Fig. 1), with *K*_*m*_ values of 16.4 ± 2.3 μM and 34.3 ± 2.2 μM, respectively. IMT M^pro^ exhibited only 6% of the catalytic efficiency compared with mature M^pro^. As previously reported, the M^pro^ N-terminal is fundamental for dimerization and any additional residues would reduce or even abolish its activity ^9,16–18^. As expected, C145S M^pro^ has only shown residual activity (Fig. 1). All three M^pro^ constructs exhibited similar thermal-stability profiles, indicating similar folding (Fig. 1).

**Fig 1.**
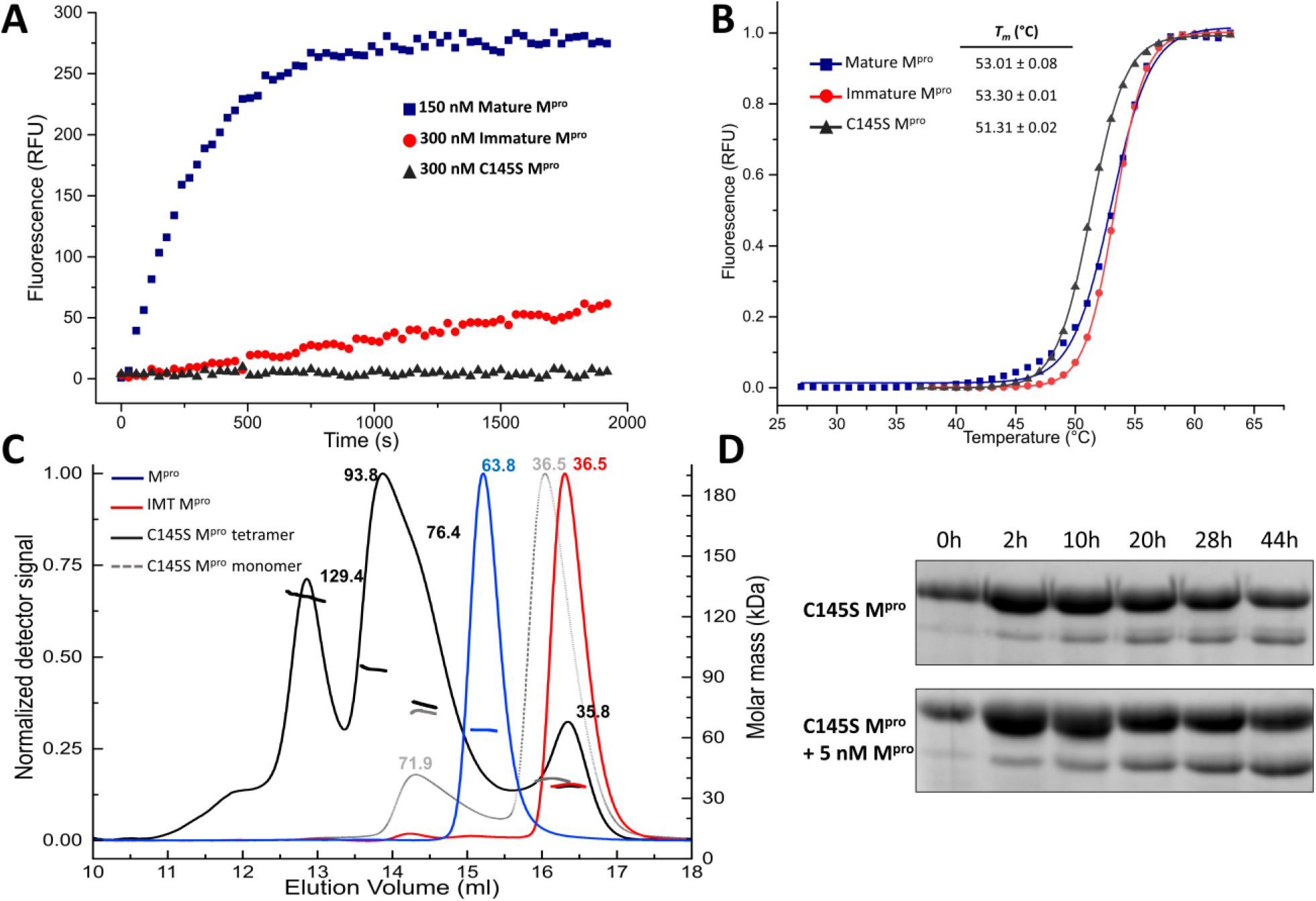
(a) Time-course reactions of M^pro^ constructs against fluorogenic peptide substrate. (b) Differential scanning fluorimetry of M^pro^ constructs. M^pro^ is shown as blue squares, IMT M^pro^ is shown as red spheres and C145S M^pro^ is shown as black triangles (c) Size exclusion chromatography elution profiles with overlaid calculated molar mass from elution peaks. M^pro^ (blue) elutes as a single peak with a calculated molecular mass consistent with a dimer. IMT M^pro^ (red) exhibits a single peak with a mass compatible with a monomer in solution. The monomeric SEC peak of C145S M^pro^ (grey) elutes as an equilibrium between dimers and monomers in solution. The tetrameric SEC peak of C145S M^pro^ (black) contains peaks of monomers, dimer, trimers and tetramers. (d) SDS-PAGE of N-terminal cleavage over time from C145S M^pro^. At top, reaction containing 10 µM C145S M^pro^, and at the bottom the same reaction supplemented with 5 nM M^pro^.

Analysis in solution using SEC-MALS suggests that M^pro^ behaves as a dimer in the tested conditions, as expected ^11^. For IMT M^pro^, the additional residues at N-terminal seem to prevent dimerization completely. For C145S M^pro^, however, the additional residues allow the protein to adopt multiple conformational states ranging from monomers to tetramers (Fig. 1).

Despite the site-direct mutagenesis of the C145S M^pro^, the enzyme exhibited residual proteolytic activity which allowed us to observe self-processing by SDS-PAGE in the course of two days. Despite the efficiency of M^pro^, its addition to the reaction does not seem to enhance significantly the rate of self-cleavage, suggesting cis-cleavage as main mechanism of N-terminal cleavage (Fig. 1).

### Crystal structure of M^pro^

M^pro^ was crystallized in several conditions and its X-ray structure was determined at 1.46 Å in *C*2_1_ space group. All 306 residues were refined at the electron density to a final *R*_work_/*R*_free_ of 0.16/0.18, with 99% of Ramachandran in favored positions (Table S2). The crystal asymmetric unit contains one monomer which could be symmetry expanded to the biological dimer, following the same pattern of the majority of known structures deposited in PDB (r.m.s.d of 0.2 Å vs PDB 5RGG, for all Cα 306). The M^pro^ protomers are formed by three domains (DI, DII and DIII), with its catalytic region located at the double-barreled DII ^9^ (Fig. S2).

### Crystal structure of IMT M^pro^

The crystal structure of IMT M^pro^ at 1.6 Å was determined using 3 merged datasets (Fig S3, S4, Table S1) in *P*2_1_2_1_2_1_ space group, with two molecules in the asymmetric unit, packed in similar shape to the known biological unit of M^pro^. The structure was refined to a final *R*_work_/*R*_free_ of 0.20/0.22, with 97% of Ramachandran in favored positions (Table S2). In the recent published structures of GM-M^pro^, both apo and ligand-complexes exhibited minor differences with the mature form ^16^. However, in our structure there are distinguishable differences in the overall structure, especially in the position of DIII helices (Fig. 2). Although IMT M^pro^ asymmetric unit resembles the biological dimer form of native protein, PISA^19^ analysis indicate that the dimer packing is unstable in solution, with an interface area of 1,256 Å^2^ (vs 1,557 Å of M^pro^), calculated free energy ΔG of −13.4 kcal/mol (vs −14.9 kcal/mol of M^pro^) for 26 potential hydrogen bonds (vs 49 of M^pro^) and 5 potential salt bridges (vs 10 of M^pro^).

**Fig 2.**
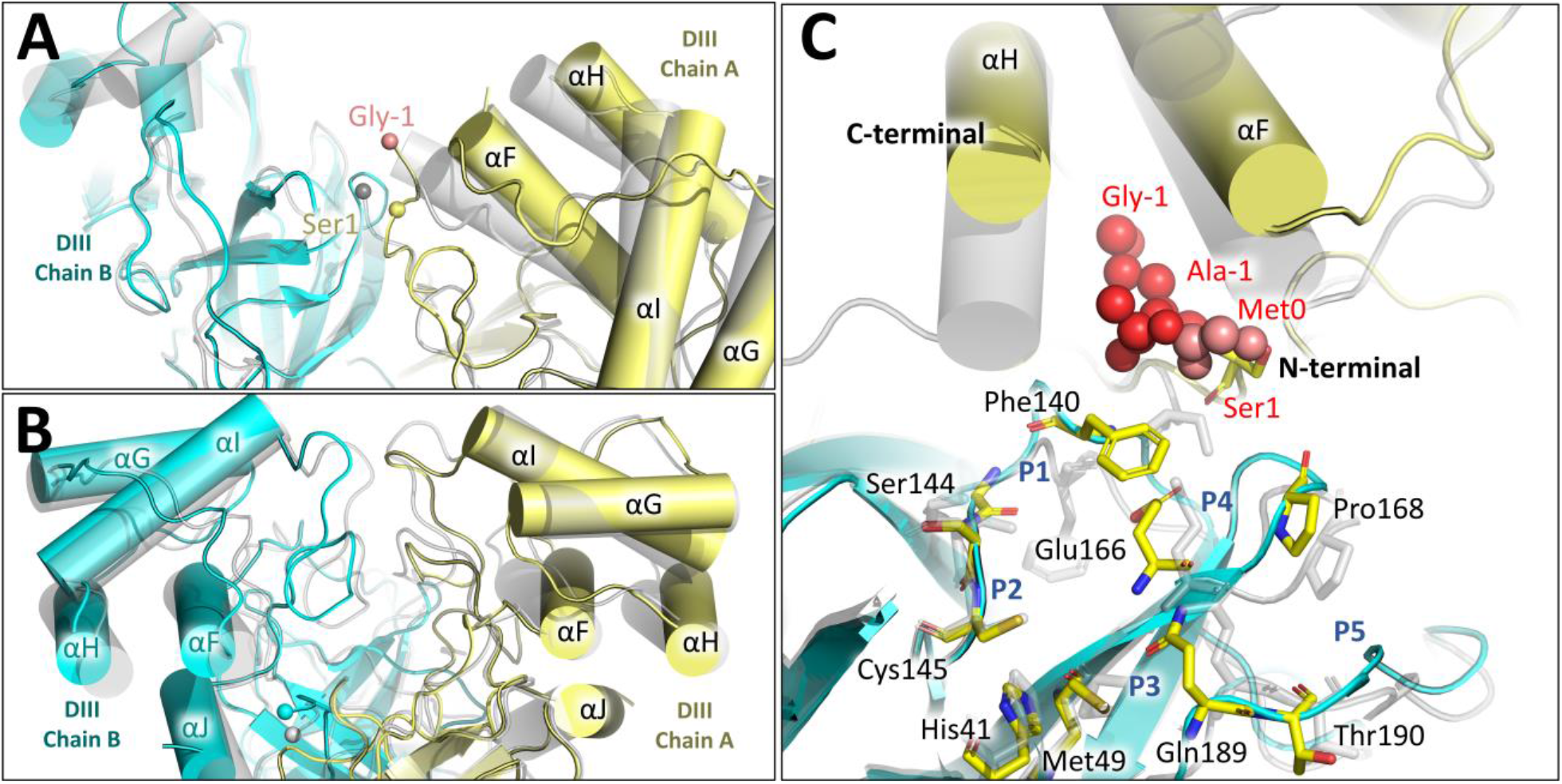
(a) Overview of DIII region from IMT M^pro^ (chain A yellow and B cyan) superposed with M^pro^ (grey ghost). N-terminal residues are depicted as spheres. (b) Rotated view showing IMT M^pro^ DIII from a distinct angle. (c) Active site residues of IMT M^pro^ chain B (cyan cartoon) superposed with M^pro^. Catalytic residues are depicted as yellow sticks. N-terminal chain A residues are depicted as spheres. M^pro^ structure and residues are shown as a grey ghost.

While IMT M^pro^ DI and DII are less affected by the N-terminal insertion (r.m.s.d of 0.34 Å vs Mpro^mat^ for Cα of 1-184), DIII appears to adopt a more open conformation relative to M^pro^ (r.m.s.d of 1.33 Å for Cα of 201-301) (Fig. 2), with the interfacing residues Ala285 at a distance of 10 Å in the IMT M^pro^ (vs 5.2 Å in M^pro^) (Fig. S5). This conformation is more accentuated at chain A where the electron density of the N-terminal insertion is clearly visible in the model. For this chain, the N-terminal insertion pushes chain A helices αF and αH further of chain B active site, opening a cleft for Phe140 rises to the surface of the molecule, leading to major conformation alterations of the chain B active site souring residues, such as Glu166, Pro168 and Gln189 (Fig. 2 and S6). The plasticity of SARS-CoV-2 M^pro^ active site was already reported when apo X-ray structures collected at cryo and room temperatures were compared ^20^, and its expected given the broad spectrum of endogenous substrates that M^pro^ is expected to process. However, the IMT M^pro^ revealed major structural alterations in the oxyanion hole, likely affecting enzyme processing. Despite the significant changes of the active site, relative position of the catalytic dyad Cys145-His41 remains unchanged in this form (Fig. 2).

### Fragment Screening of M^pro^ immature

Recently, a small-fragment library of more than 1,250 unique fragments were screened against SARS-CoV-2 M^pro^, identifying 74 high-value fragment hits, including 23 non-covalent and 48 covalent hits in the active site, and 3 hits close to the dimerization interface ^21^. In here, we applied the same technique to probe new druggable cavities in IMT M^pro^. Although the difference in scale of our experiment, we were able to identify five distinguishable sites in this form of the protein (Fig. 3). Site #1 is the active site of chain A, in which fragment f2xe03 was identified interacting with Glu166 N and Cys145 S. Interesting, a unique cavity marked as Site #3 was identified in our experiments, bound to fragment f2xg02 by Arg4 main chain O. That cavity lies between the interface of chains A and B, and is not present in M^pro^ which adopt a more closed conformation. This new site and fragment could serve as an anchor for development of new inhibitors targeting M^pro^ dimerization process, a mode of action that was too date only theorized ^22^. Details about data processing and statistics are given in Table S3.

**Fig 3.**
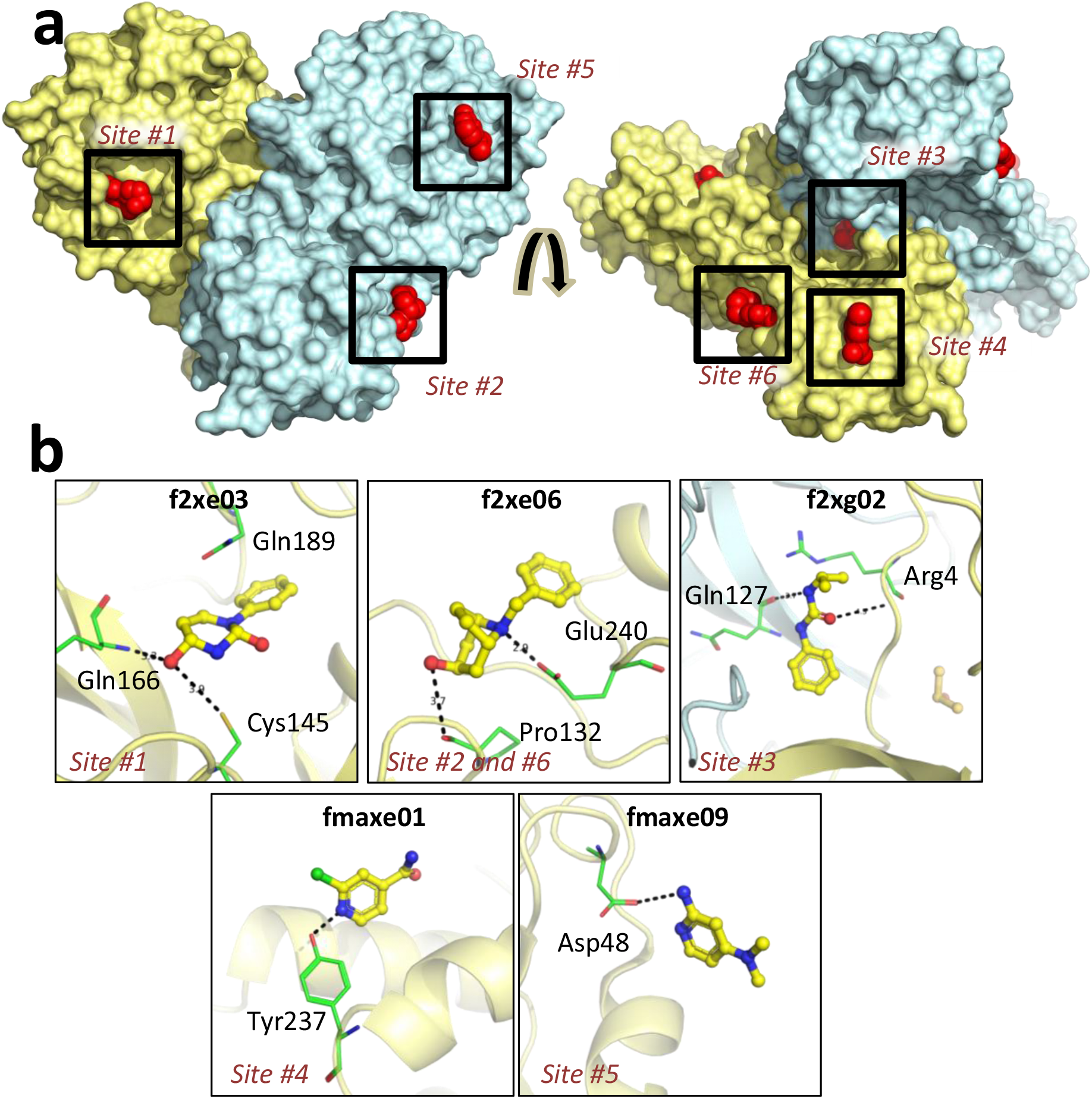
(a) Location of IMT M^pro^ probing fragments identified during screening. Chain A is colored as yellow surface, chain B as cyan surface. Fragments are depicted as red spheres. (b) Contact details of identified fragments. Chain A is colored as yellow cartoon and chain B as cyan cartoon. Fragments are depicted as yellow sticks. Residues forming polar contacts are depicted as green lines. Contacts are depicted as black dashes.

### Crystal structure of C145S M^pro^ in complex with N and C-terminal residues

The tetramer peaks were crystallized and X-ray structure determined at 2.8 Å and *R*_work_/*R*_free_ of 0.20/0.24 (Table S2), revealing a new crystal form in which N-terminal chain B residues are trimmed in the active site of chain A, occupying subsites S1-S5 (Fig. 4). Despite the site directed mutagenesis of the catalytic cysteine to serine, electron density shows that Gln0 and Ser1 are non-covalently bound in the amino region, clearly indicating that the N-terminal cis-cleavage was completed. At the S1 subsite, Gln0 NE2 interacts with Glu166 OE1 by a hydrogen bound (2.7 Å), while Gln0 form interacts with Ser145 in the position of the native oxyanion hole (Fig. 4, Fig S8-S9). To accommodate the hydrophobic sidechain of Leu-1 at P2, Met49 and Met165 are pushed further of each other (Fig. S10), leading to a more opened groove of this subsite relatively to the apo-state, explaining the ability of this subsite to accommodate a variety of hydrophobic side chain residues, such as Leu, Met, Ile, Val and Phe ^6,23^. Yet, from the eleven endogenous recognition sites of coronaviruses, S2 Leu carrying sequences are the ones in which M^pro^ display higher catalytic efficiency, highlighting the importance of this conformation for drug design. At subsites S3-S5, the interactions of Val-2, Ala-3 and Ser-4 are mainly maintained by hydrogen bounds between the polar residues of protein and peptide side chains (Fig. 4), which explains the ability of M^pro^ to recognize a large variety of chain sequences at those positions.

**Fig 4.**
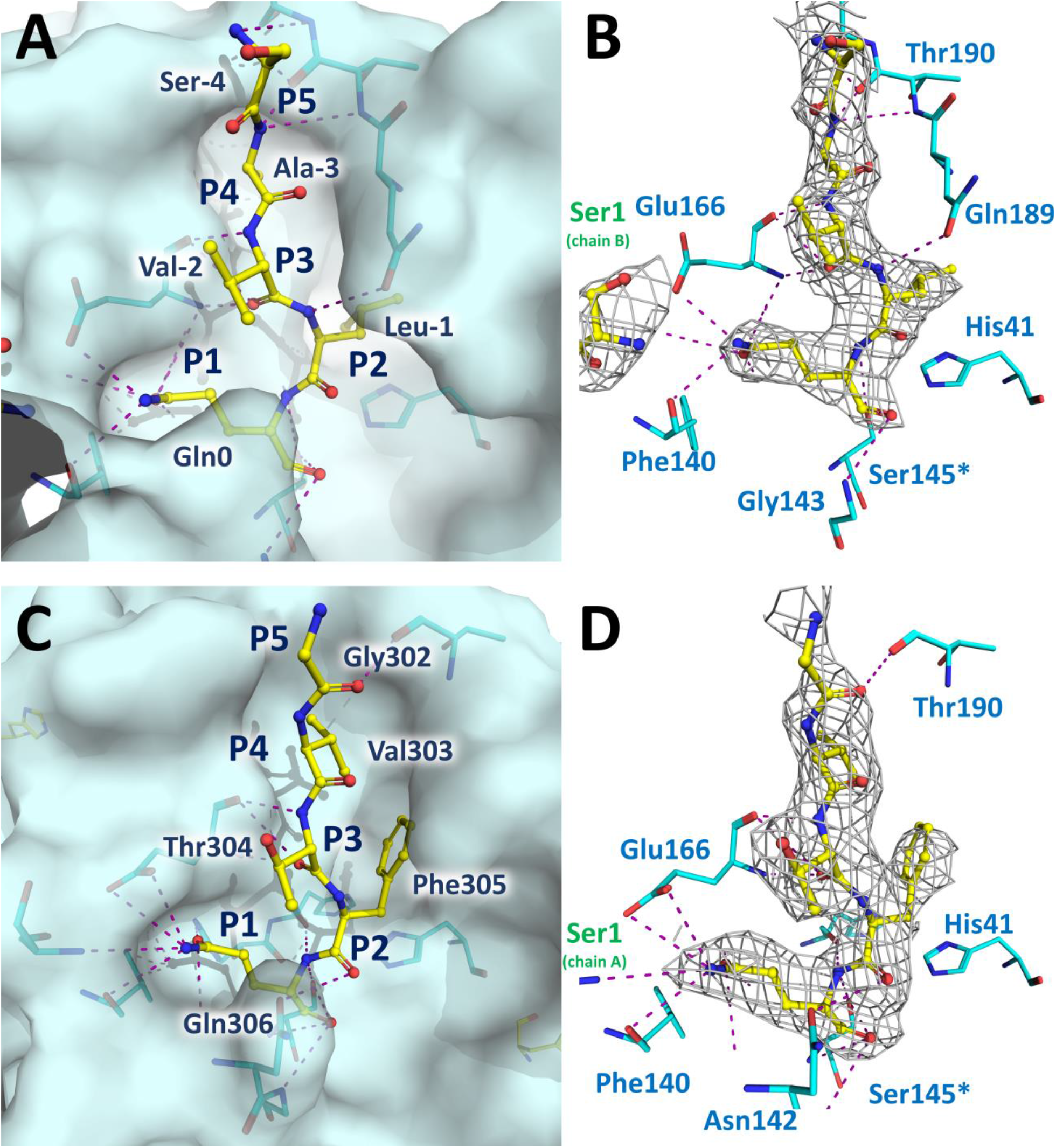
(a) C145S M^pro^ chain A active site (cyan surface) in complex with processed N-terminal residues (yellow sticks). Main interacting residues are depicted as blue lines. (b) C-terminal peptide (yellow) main interactions with C145S M^pro^ chain A active site residues (blue). (c) C145S M^pro^ chain B active site (blue surface) in complex with processed C-terminal residues (yellow sticks). Main interacting residues are depicted as blue lines. (d) C-terminal peptide (yellow) main interactions with C145S M^pro^ chain B active site residues (blue). For (b) and (d), the 2mFo-DFc electron density contoured at 0.8σ. Ser1 from respective dimerization partners are depicted with green letters. *Ser145 is the site-direct mutant of Cys145. Simulated annealing omit map is available in Fig S7.

The crystal structure of C145S M^pro^ revealed another important step in the maturation process of M^pro^. At the same time that chain B N-terminal additional residues are processed by chain A, its C-terminal residues (301-306) are almost 180° twisted from its original position (Fig. S11) and trimmed in the active site of a symmetric related chain B (Fig. 4) a phenomenon that was also recently described by another group in both native and in a C145A M^pro^ mutant ^24^. During this event, two C145A M^pro^ dimers appear to be linked by the interaction of the C-terminal and a respective active site, assuming a tetrameric conformation (Fig. 5). Within the active site, Gln306 occupies the respective position of Gln0 at S1, while S2 is occupied by Phe305, increasing the distance between Met49 and Met165 relatively to chain A bound to N-terminal (Fig. S10). As the N-terminal residues, subsites S3-S5 interactions with C-terminal are mainly maintained by hydrogen bounds between main chains (Fig 4).

**Fig 5.**
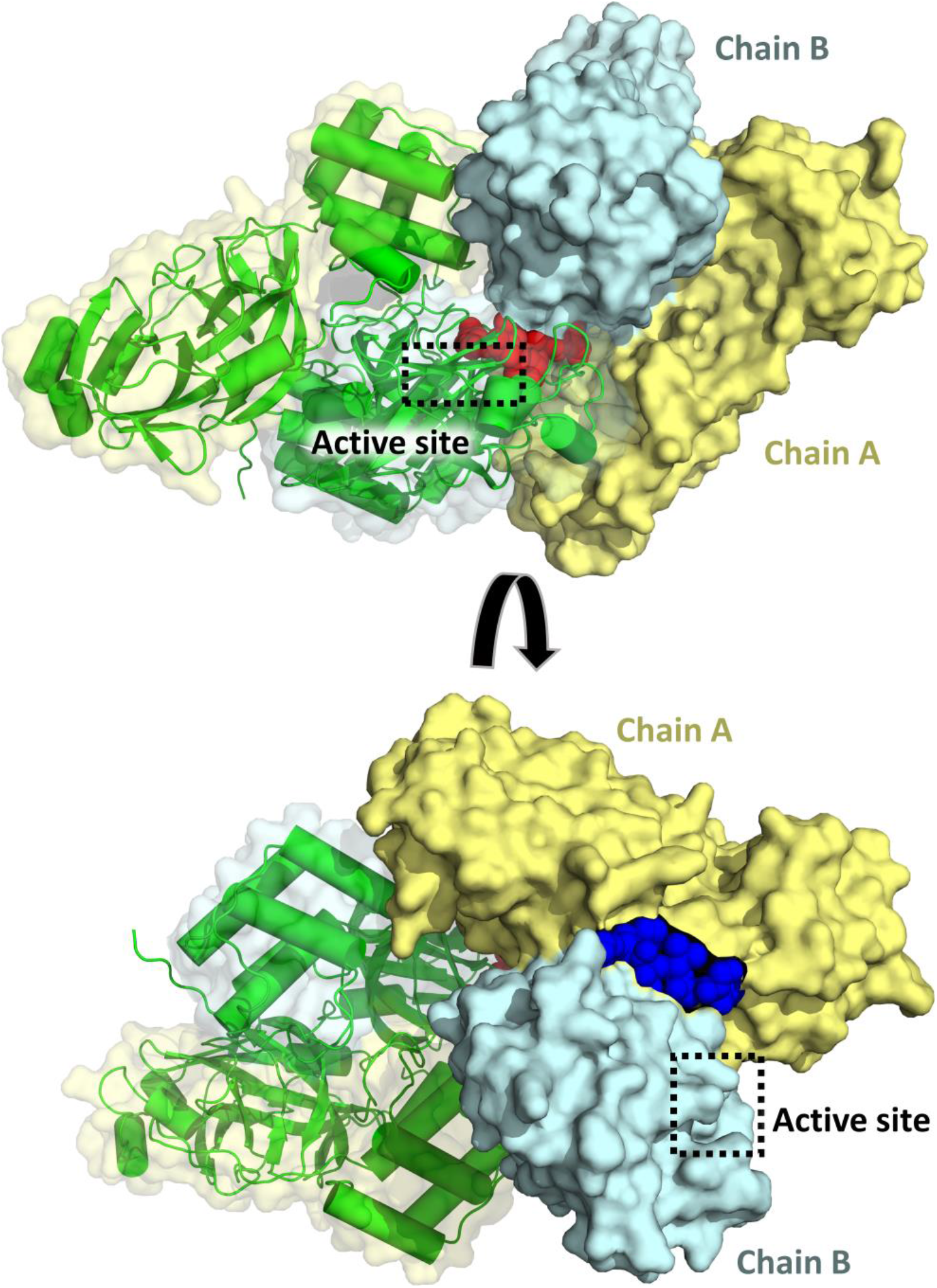
Overview of the tetrameric intermediary formed by C145S M^pro^ during self-processing. Chain A is colored as yellow surface, chain B as cyan surface. *Trans*-cleavage M^pro^ partner is show as green cartoon. N-terminal residues are depicted as blue spheres, and C-terminal residues are depicted as red spheres.

### The maturation process of M^pro^ and its impact on Drug discovery

M^pro^ is firstly produced as the Nsp5 domain of the viral polyproteins before they are proteolytically processed into 15 or 16 non-structural proteins ^12^. Immediately after translation, the immature form of M^pro^ would contain both N and C-terminal insertions that need self-processing to generate the mature form of the enzyme ^13^.

The biochemical assays of IMT M^pro^ revealed that a minor insertion at the N-terminal produces a protein form that behaves as a monomer in solution and its almost depleted of enzymatic activity, even though the general folding remains similar to the full mature form. The same process occurs to C145S M^pro^ with native N-terminal inserted residues, although, in this case, a slow cleavage of the N-terminus results in the formation of dimers overtime (Fig. 1). It is important to highlight that when M^pro^ is added to C145S M^pro^, the N-terminal cleavage and dimer formation does not seem to be enhanced, strongly suggesting that this initial maturation step is a *cis*-cleavage event. This is in agreement with the model proposed by Li and colleagues (2010) in which two M^pro^ form a transient dimer that is stabilized by the binding the N-terminal site of its substrate (another M^pro^ in polyprotein) and further cleave to free its N terminus ^14^.

After the active site region is matured (or even concomitantly), dimeric M^pro^ C-terminal seem to assume an unusual twisted position (Fig. S11), allowing it to be docked into the active site of another mature or half-mature M^pro^ dimer (Fig. 4). In this step, the trans-cleavage processing of the C-terminal residues would serve as an anchor for a transitory tetrameric state of the protein, herein captured with the construct of the mutant C145S M^pro^ with the processing of the N-terminal residues (Fig. 5). As a result, full mature M^pro^ is produced and its ready to process other parts of the viral polyprotein. During all those maturation processes, both M^pro^ active site and surface undergo significant conformational changes, which could guide targeted drug development (Fig. 6 and Scheme S1). Our results not only shed light in the self-maturation process of SARS-CoV-2 M^pro^, but also bring the perspective of developing drugs targeting intermediate states of this enzyme.

**Fig 6.**
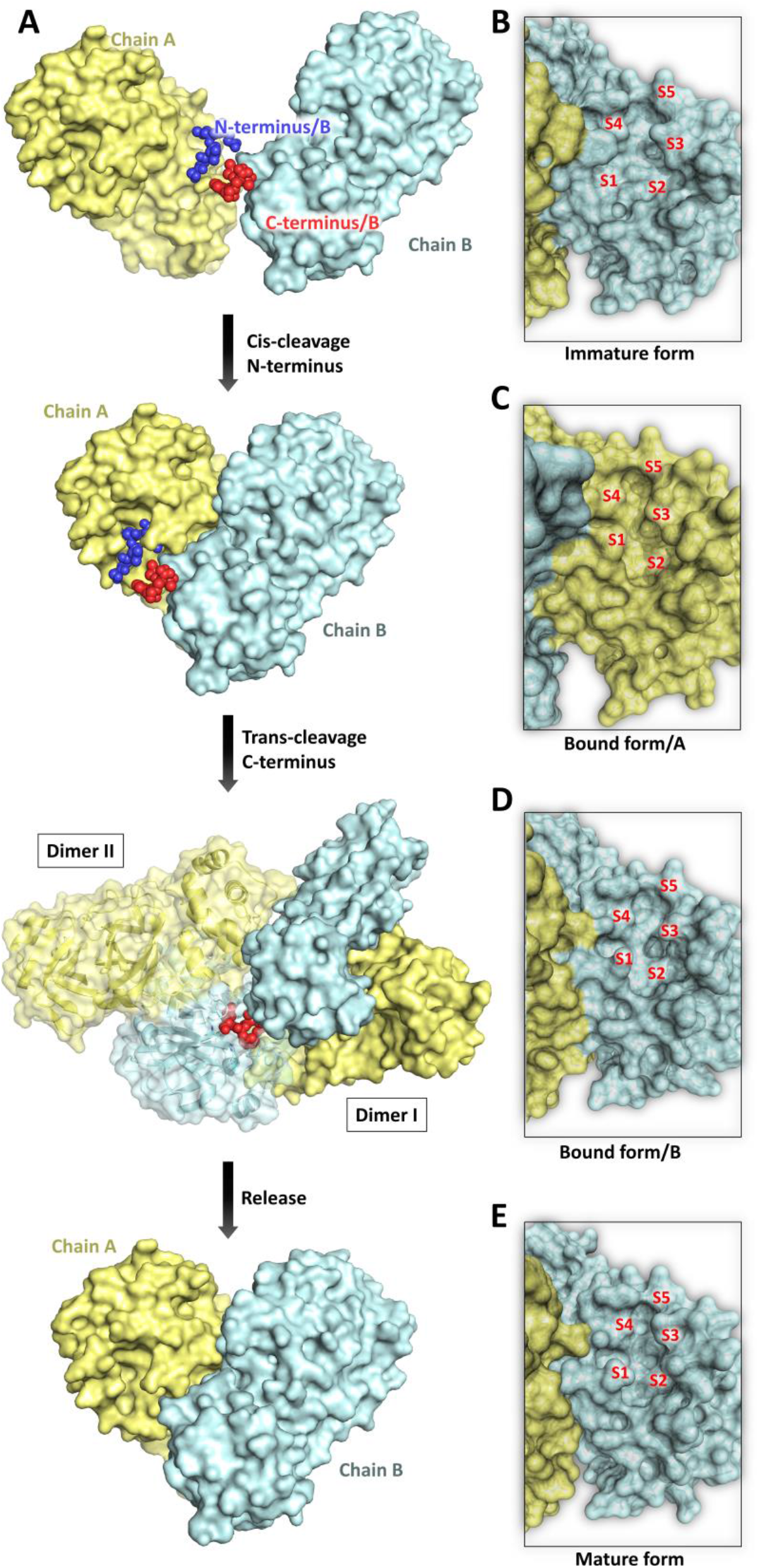
(a) Scheme containing steps of SARS-CoV-2 M^pro^ self-maturation process. At first, two protomers assembly as an immature dimer during N-terminal cis-cleavage. After processing, M^pro^ has a mature active site, which allows the stabilization of the dimeric form. The dimer C-terminal is them trans-cleaved by another full or half mature dimer, producing the full mature form of M^pro^. (b) Surface view of chain B active site from immature form. (c) Surface view of chain A active site during N-terminal residues recognition. (d) Surface view of chain B active site during C-terminal residues recognition. (e) Surface view of full mature M^pro^ active site.

## Acknowledgments

Authors acknowledge SIRUS (Campinas, Brazil) proposal 20200014. F2X Entry was obtained with resources from Federal Ministry of Education and Research (BMBF), while FragMAXlib with resources from Swedish Research Council (VR).

## Author contributions

ASG conceived this project. GDN performed biochemical and biophysical experiments. GDN, ASG, AMN, VOG and GMAL performed crystallographic experiments and analysis. ASG, GDN, AMN, VOG, GO and RSF wrote this manuscript. HVDR performed SEC-MALS experiments. Other authors supported experiments and data analysis.

## Data and materials availability

Structure factors and atomic coordinates have been deposited with the protein data bank with accession codes PDB ID 7KFI, 7KPH, 7KVG, 7LFE, 7LDX, 7LFP, 7KVL and 7KVR. Other data are available from the corresponding author upon reasonable request.

## Supplementary Materials

### Cloning and expression of SARS-CoV-2 M^pro^ forms

The viral cDNA template (GenBank MT126808.1), kindly provided by Dr. Edison Durigon (University of São Paulo, São Paulo, Brazil), was synthetized using the SCRIPT One-Step RT-PCR kit (Cellco Biotec) and random hexamers primers. For production of IMT M^pro^, coding region of M^pro^ (residues 3264-3569) was amplified using primers: Fw 5’ CAGGGCGCCATGAGTGGTTTTAGAAAAATGGCATTC 3’ and Rv 5’ GACCCGACGCGGTTATTGGAAAGTAACACCTGAGAC 3’, and the sequence was inserted into the pET_M11 vector, which encodes an N-terminal 6xHis-tag followed by a TEV protease cleavage site (ENLYFQ↓GAM), using the LIC method (*25*), forming the plasmid pET_M11-IMT-M^pro^. To obtain the mature form of M^pro^, native N-terminal residues (GAMSAVLQ↓SGFRK) were inserted into pET_M11-IMT-M^pro^ by inverse PCR using primers: Fw: 5’ GCTGCAGAGTGGTTTTAGAAAAATGGCATTC 3’ and Rv: 5’ ACGGCTGACATGGCGCCCTGAAAATA 3’. Amplified product was treated with T4 Polynucleotide Kinase (PNK, Thermo Fischer Scientific) and T4 Ligase (Cellco Biotec), forming plasmid pET_M11-M^pro^. For C145S M^pro^ construct, pET_M11-M^pro^ was used as template for inverse PCR with primers Fw 5’ CCTTAATGGTTCATCTGGTAGTG 3’ and Rv 5’ AATGAACCCTTAATAGTGAAATTGG 3’. The PCR product was digested with DpnI (NEB), followed by treatment with PNK and T4 DNA ligase, forming the pET_M11-C145S-M^pro^. All plasmids were transformed in DH5α *E. coli* competent cells. All PCRs were conducted with FastPol (Cellco Biotech). Positive clones were selected and confirmed by sequencing. Schematics of constructs are given in Fig. S1.

For protein production, *E. coli* BL21 cells were transformed with respective plasmids and cultured in ZYM-5052(*26*) at 37^°^C and 200 RPM to an OD_600_ of 0.8, followed by expression at 18 °C, 200 RPM for 16 h. Cells were harvested by centrifugation at 5,000 x *g* for 40 min at 4 °C, resuspended in lysis buffer (20 mM Tris pH 7.8, 150 mM NaCl, 1 mM DTT), disrupted by sonication and the lysate was clarified by centrifugation at 12,000 x *g* for 30 min at 4 °C.

### Protein purification of IMT M^pro^

After expression, a large amount of IMT M^pro^ had its 6xHis-tag cleaved by autoproteolytic process. The small fraction of 6xHis tagged protein was removed from the lysate using Ni-NTA resin (Qiagen). The cleaved protein was purified by adding 1 M ammonium sulfate to the cell lysate followed by incubation on ice for 10 min. The precipitated protein was recovered by centrifugation at 12,000 x *g* for 30 min at 4 °C, resuspended in gel filtration buffer (20 mM Tris pH 7.8, 150 mM NaCl, 1 mM EDTA, 1 mM DTT) and purified by size-exclusion chromatography using a HiLoad 26/100 Superdex 200 column (GE Healthcare) pre-equilibrated with gel filtration buffer. Purified fractions were aliquoted, flash-frozen and stored at −80 °C for enzymatic assays and crystallization. For crystallization, protein was concentrated to 14 mg.mL^-1^ using 10 kDa MWCO centrifugal concentrators (Vivaspin, Sartorius). Protein concentrations were determined using the measured absorbances at 280 nm and the theorical extinction coefficient of 32,890 M^-1^.cm^-1^. Protein purity was analyzed by SDS-PAGE (Fig. S1).

### Protein purification of M^pro^

M^pro^ was purified similar to IMT M^pro^, with an additional step of cation exclusion chromatography. After the size exclusion chromatography, the protein was buffer exchanged to 20 mM Tris pH 8.0, 1 mM DTT, and then injected into a Mono-Q 5/50 GL column (GE Healthcare). Protein was eluted using a linear gradient of a buffer containing 20 mM Tris pH 8.0, 1 M NaCl and 1 mM DTT. Finally, fractions containing the purified protein were buffer exchanged to gel filtration buffer. Purified fractions were aliquoted and protein was concentrated and quantified similarly to IMT M^pro^ (Fig. S1).

### Protein purification of C145S M^pro^

For C145S M^pro^, protein was purified by immobilized metal chromatography (IMAC) using a 5 mL HisTrap FF column (GE Healthcare). After column washing with buffer A (20 mM Tris pH 7.8, 150 mM NaCl, 25 mM Imidazole), protein was eluted with buffer A supplemented with 250 mM imidazole. Sample was buffer exchanged using a 5 mL HiTrap desalting column (GE Healthcare) equilibrated with buffer A. To remove the 6xHis-tag, 2 mg of TEV protease and 4 mM DTT were added to the sample and incubated for 16 h at 4 °C. Next day, non-cleaved protein and TEV were removed by a second step of IMAC in buffer A. Finally, the protein was purified by size-exclusion chromatography using a HiLoad 16/60 Superdex 75 column (GE Healthcare) equilibrated with gel filtration buffer. Purified fractions were aliquoted, and protein was concentrated and quantified as described for other constructions (Fig. S1).

### Crystallization

Crystallization screening was performed with the sitting drop vapor diffusion method in 96 -well plates using a Phoenix Liquid Handling System-Gryphon LCP (Art Robbins Instruments) and commercially available kits at 20 °C. For M^pro^, crystals appeared after 1 day in 0.1 M Bis-Tris pH 6.5, 25% PEG 3,350, which were cryo-protected using the reservoir solution and 30% PEG 400. Crystals of IMT M^pro^ were observed in several conditions. After optimization, crystals grown in 0.1 M MES pH 6.5, 10% 2-propanol, 20% PEG Smear Low (BCS Screen, Molecular Dimensions) were used as seeds for the diffraction crystals, obtained in 0.1 M MES pH 6.7, 5% DMSO, 8% PEG 4,000 (*21*). Crystals of C145S M^pro^ were obtained after 3 days in 0.1 M phosphate/citrate, pH 5.5, 20% v/v PEG Smear High (BCS Screen, Molecular Dimensions).

### Data collection and processing at MANACA Beamline

During the initial commissioning phase (July to October 2020) the MANACA (MAcromolecular Micro and NAno CrystAllography)(*27*) beamline adopted an emergency commissioning plan to deliver the basic instrumentation to perform data collection of SARS-CoV-2 related samples. Thus, during this period, the beamline has operated on a fixed-energy regime (9 keV) with manual crystal mounting, single-axis goniometry, beam flux estimated to be about 1·10^11^ ph/s/10 mA at 9.15 keV and adjustable beam size from about 18 (h) x 20 (v) µm^2^ to 100 (h) x 80 (v) µm^2^ (FWHM). This project was the first external user session at MANACA beamline and the first operating beamline at Sirius (SIRIUS, Brazil). The focus was optimised to 61 (h) x 48 (v) µm^2^ at sample position (Fig. S3). Even without the full capabilities, the beamline opening was very important to SARS-CoV-2 structural biology studies.

X-ray data for apo IMT M^pro^ was collected from three isomorphous independent crystals, that were processed by XDS via autoPROC (*28, 29*). Data herein was used for confirm reliability of the beamline (Fig. S5 and Table S1). Datasets were then scaled and merged using Aimless (*30*), and the resulting dataset was used for structural determination of IMT M^pro^ by molecular replacement with Phaser (*31*) using PDB 5RGQ as template. Model was refined with COOT(*32*) and BUSTER (*33*) at 1.6 Å and deposited under the code 7KFI.

X-ray data for mature M^pro^ and C145S M^pro^ were processed by XDS via autoPROC (*28, 29*) and scaled using Aimless (*30*). Mature M^pro^ and C145S M^pro^ were solved by molecular replacement with Phaser (*31*) using template models 5RGQ and 7KFI, respectively. Mature M^pro^ and C145S M^pro^ were refined with COOT(*32*) and phenix.refine (*34*), and are respectively deposited under the codes 7KPH (at 1.4 Å) and 7KVG (at 2.8 Å). Details of data processing parameters and statistics are given in Table S2.

### Fragment Screening of IMT M^pro^

For the fragment screening of IMT M^pro^, we used the settled plates fragment libraries of FragMAXlib (Talibov *et al*., to be published) and F2XEntry (*35, 36*). In those plates, the content of each drop-well was resuspended in 1.0 µL of 0.1 M MES pH 6.7, 5% DMSO (v/v), 8% PEG 4,000 (w/v), 30% PEG 400 (w/v), and crystals were added afterwards. After 4 h soaking at room temperature, crystals were manually harvested and flash cooled for data collection.

During the commissioning phase of MANACA, 166 of those crystals were tested, leading to 77 usable datasets. To analyze the data, a simplified version of FragMAXapp was configurated in our laboratory end-station computer. Within FragMAXapp, restrictions libraries were generated by phenix.eLBOW (*37*) using rm1 force field for geometry optimization, and datasets were processed through autoPROC/STARANISO or DIALS via XIA2 (*28, 38, 39*). Molecular replacement and initial refinement were performed using DIMPLE (*40*) using PDB 7KFI as template. To highlight electron density of weak binding events, map averaging and statistical modelling were performed by PanDDA software (*41*). Models were refined with COOT(*32*) and phenix.refine (*34*). Details of data processing and refinement statistics are given in Table S3.

#### Activity assays

All enzymatic assays were carried out using FRET-based substrate DABCYL-KTSAVLQ↓SGFRKM-E(EDANS)-NH_2_ in assay buffer (20 mM Tris pH 7.3, 1 mM EDTA, 1 mM DTT). M^pro^, IMT M^pro^ and C145S M^pro^ assays were performed at final concentration of 0.14 μM, 0.3 µM and 0.3 µM, respectively. Prior to reactions, enzymes were incubated in assay buffer at 37 °C for 10 min, followed by substrate addition. To determine the kinetics parameters (*K*_*m*_, *V*_*max*_ and k_cat_), the substrate was diluted to a range of concentrations from 100 μM to 0.78 μM. Initial velocity was derived from the slope of linear phase of each time-curse reaction, and Michaelis-Menten fitting was obtained using Origin Pro 9.5.1 Software (OriginLab). Fluorescence measures were performed in SpectraMax Gemini EM Microplate Reader with λ_exc_/λ_emi_ of 360/460 nm, every 30 s over 60 min at 37° C. All assays were performed in triplicates.

To test C145S M^pro^ auto-cleavage activity, 6xHis-tagged C145S M^pro^ was buffer exchanged in 20 mM Hepes pH 7.3, 100 mM NaCl, 1 mM DTT. Two reactions were prepared for comparison, one containing 10 µM C145S M^pro^ and other containing 10 µM C145S M^pro^ and 5 nM of mature M^pro^. Aliquots of each reactions were collected for the period of 40h. Samples were analyzed by SDS-PAGE on a 12.5% SDS polyacrylamide gel.

### Differential Scanning Fluorimetry

For differential scanning fluorimetry assays (DSF), SYPRO Orange at 5X final concentration was added to protein diluted to 25 μM protein in gel filtration buffer. Denaturation curves was obtained ranging temperature from 25°C to 75°C increasing one degree per cycle and fluorescence was measured in the end of each cycle. Experiment was conducted in a qPCR system Mx3000P (Agilent). The melting temperature was obtained using the Boltzmann fitting on Origin Pro 9.5.1 Software (OriginLab).

### In solution oligomeric state of M^pro^ constructs

The in solution oligomeric states of the purified samples were evaluated by size exclusion chromatography coupled with multi-angle light scattering (SEC-MALS) in running buffer composed by 20 mM Tris-HCl pH 7.8 and 100 mM NaCl. For that, 50 µL of each M^pro^ construct at concentration of 10 µM were injected in a Waters 600 HPLC system (Waters) coupled in-line with an UV detector, a miniDAWN TREOS multi-angle light scattering apparatus (Wyatt Technology), a column Superdex 200 Increase 10/300 GL (GE Healthcare), and a refractive index detector Optilab T-rEX (Wyatt Technology). The light-scattering detectors were normalized with bovine serum albumin (Sigma-Aldrich). Data were collected and analyzed with the ASTRA 7 integrated software provided by Wyatt. The flow rate used was 0.5 mL/min. Results are summarized in Table S4.

**Fig S1.**
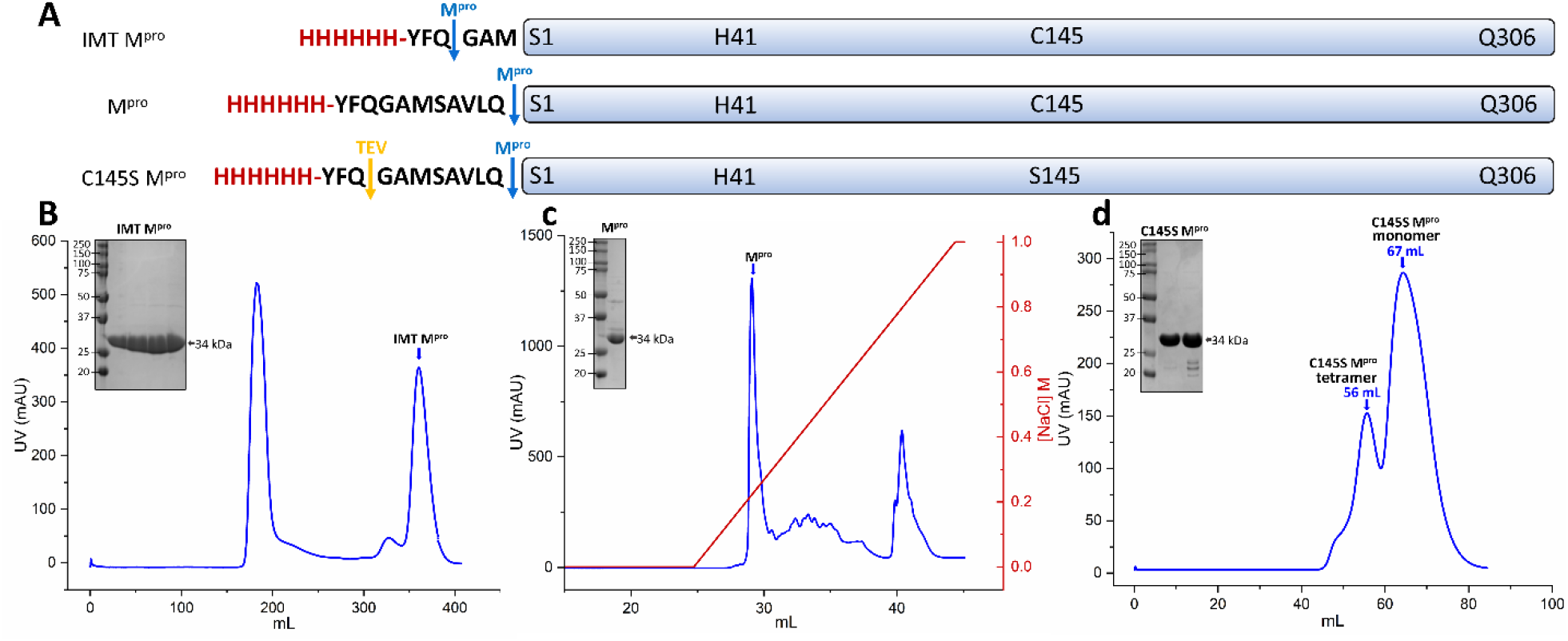
(A) Schematic showing different constructs of M^pro^. (B) Gel-filtration profile and SDS-page of purified M^pro^ constructs.

**Fig S2.**
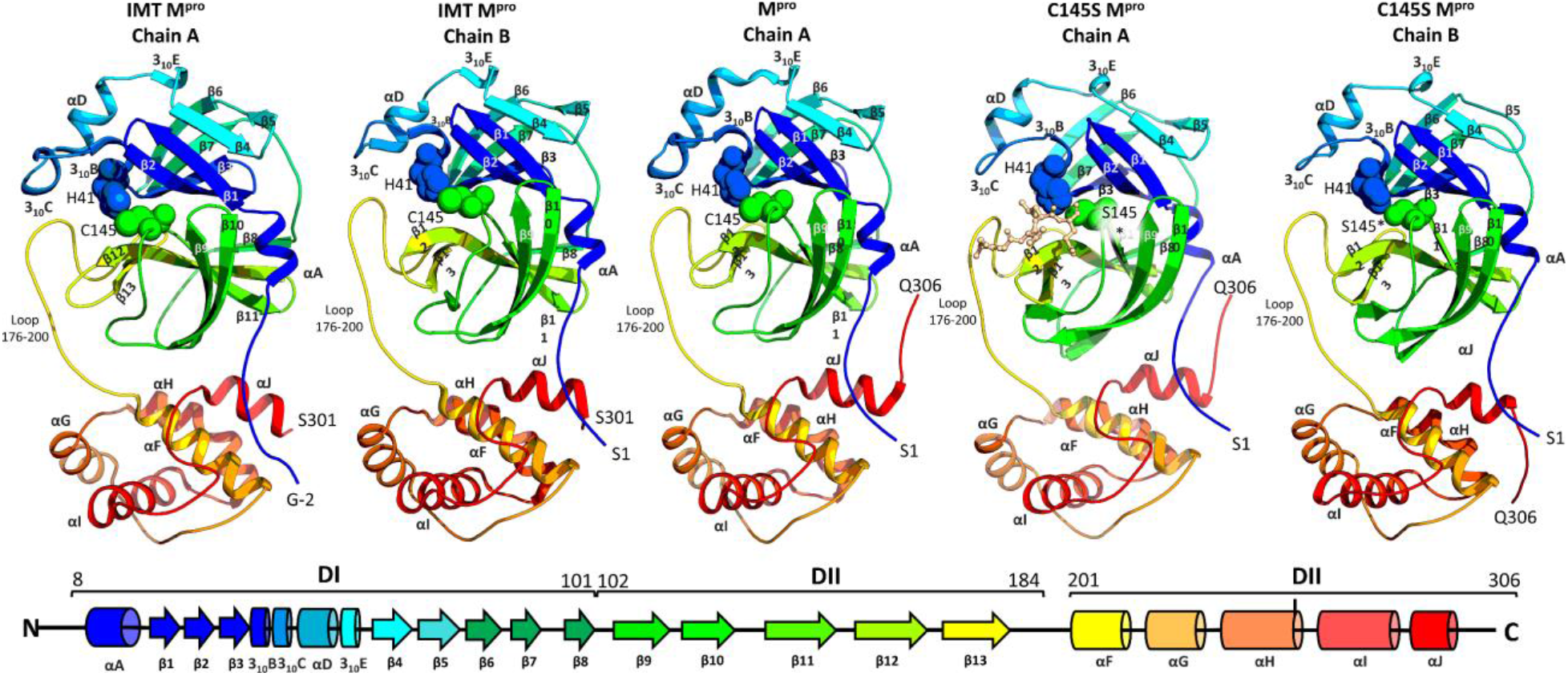
Cartoon topology and secondary structure nomenclature of each M^pro^ described in this manuscript. At bottom, alpha-helixes are draw as cylinders and beta-strands as arrows.

**Fig S3.**
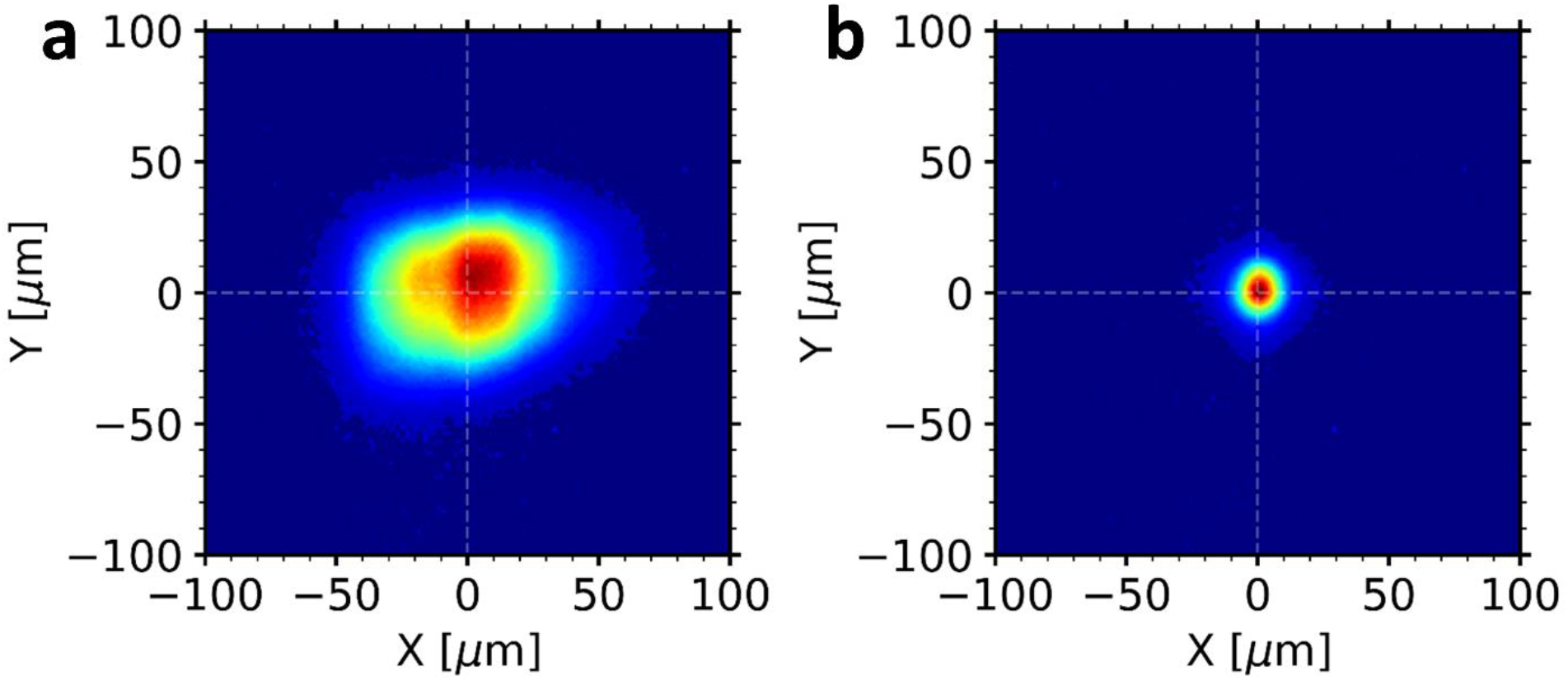
**(A)**, Unfocused beam with approximately 61 (h) x 48 (v) µm^2^ (FWHM) selected by the user to collected the data presented in this paper. **(B)**, smallest beam size reached so far during the commissioning phase, 14 (h) x 17 (v) µm^2^ (FWHM) which is very close to the nominal one (10 (h) x 7 (v) µm^2^). FWHM, full width at half maximum.

**Fig S4.**
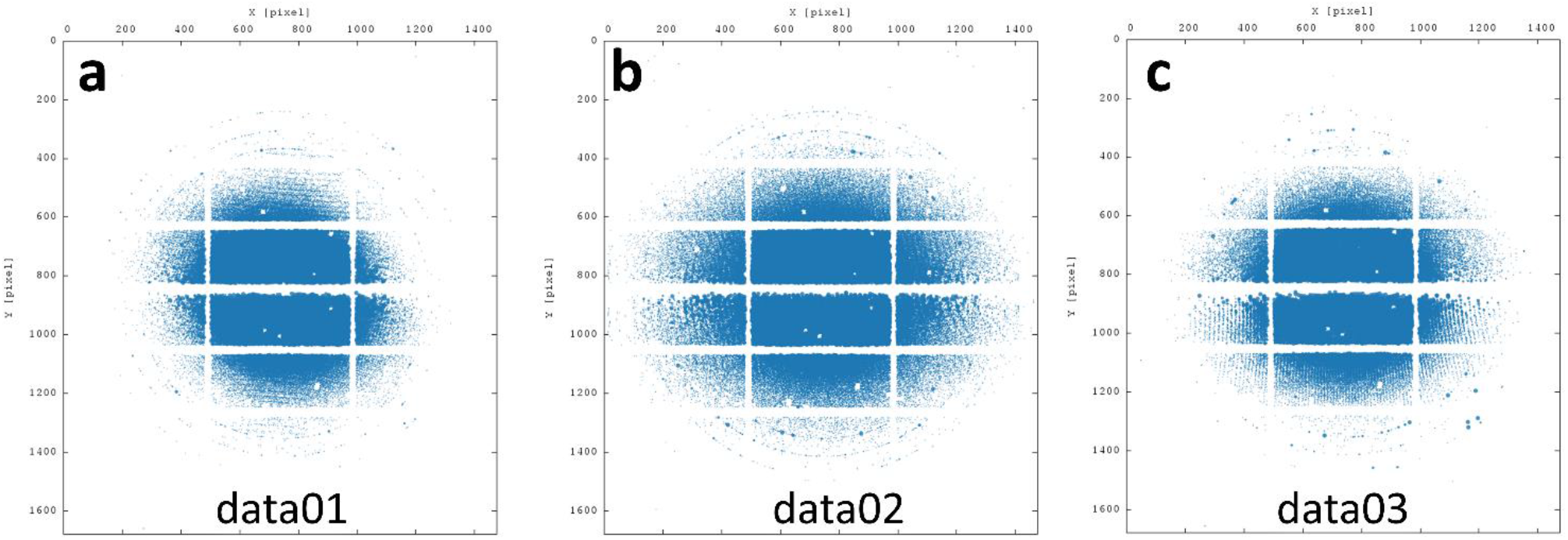
Indexed spots of IMT M^pro^ test datasets at MANACA beamline. Figures were generated with autoPROC.

**Fig S5.**
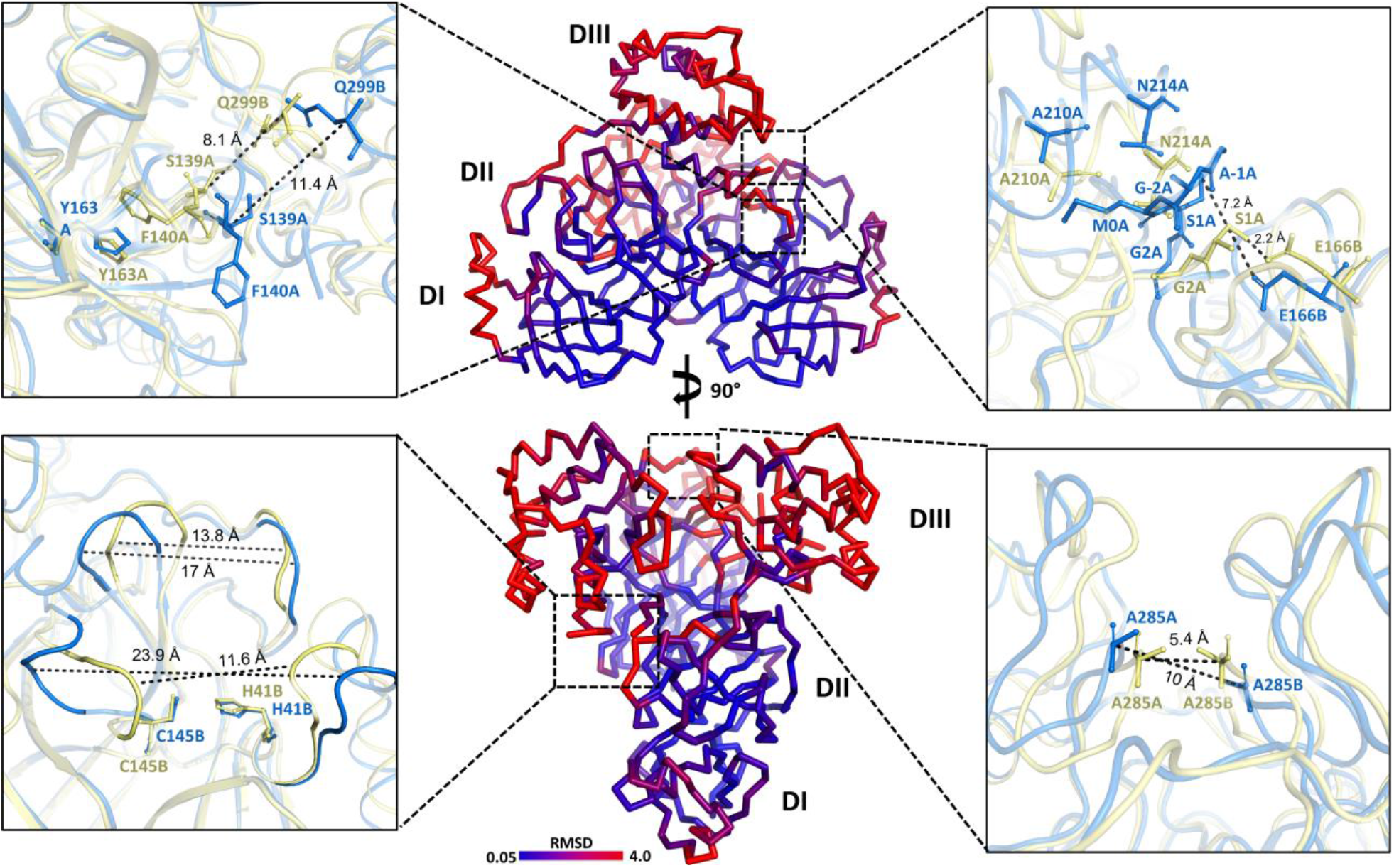
At center, superposition of M^pro^ and IMT M^pro^ dimers colored according its RMSD. At sidelines, detailed view of specific selected regions and residues, with M^pro^ colored in yellow ^and^ IMT M^pro^ colored in blue.

**Fig S6.**
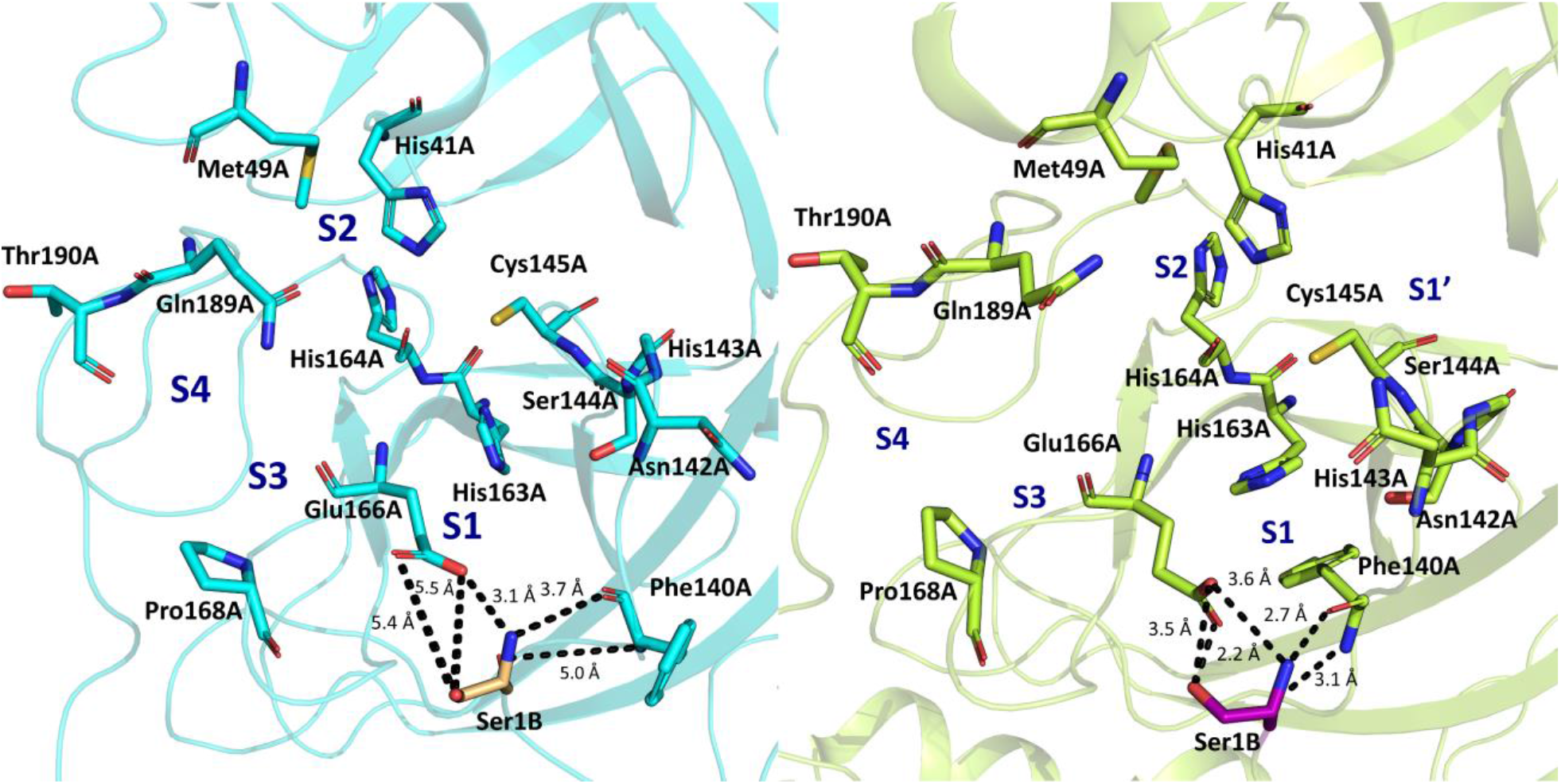
Active site of M^pro^ and IMT M^pro^. At left, IMT M^pro^ catalytic residues depicted blue sticks. At right, M^pro^ catalytic residues depicted as lime sticks.

**Fig S7.**
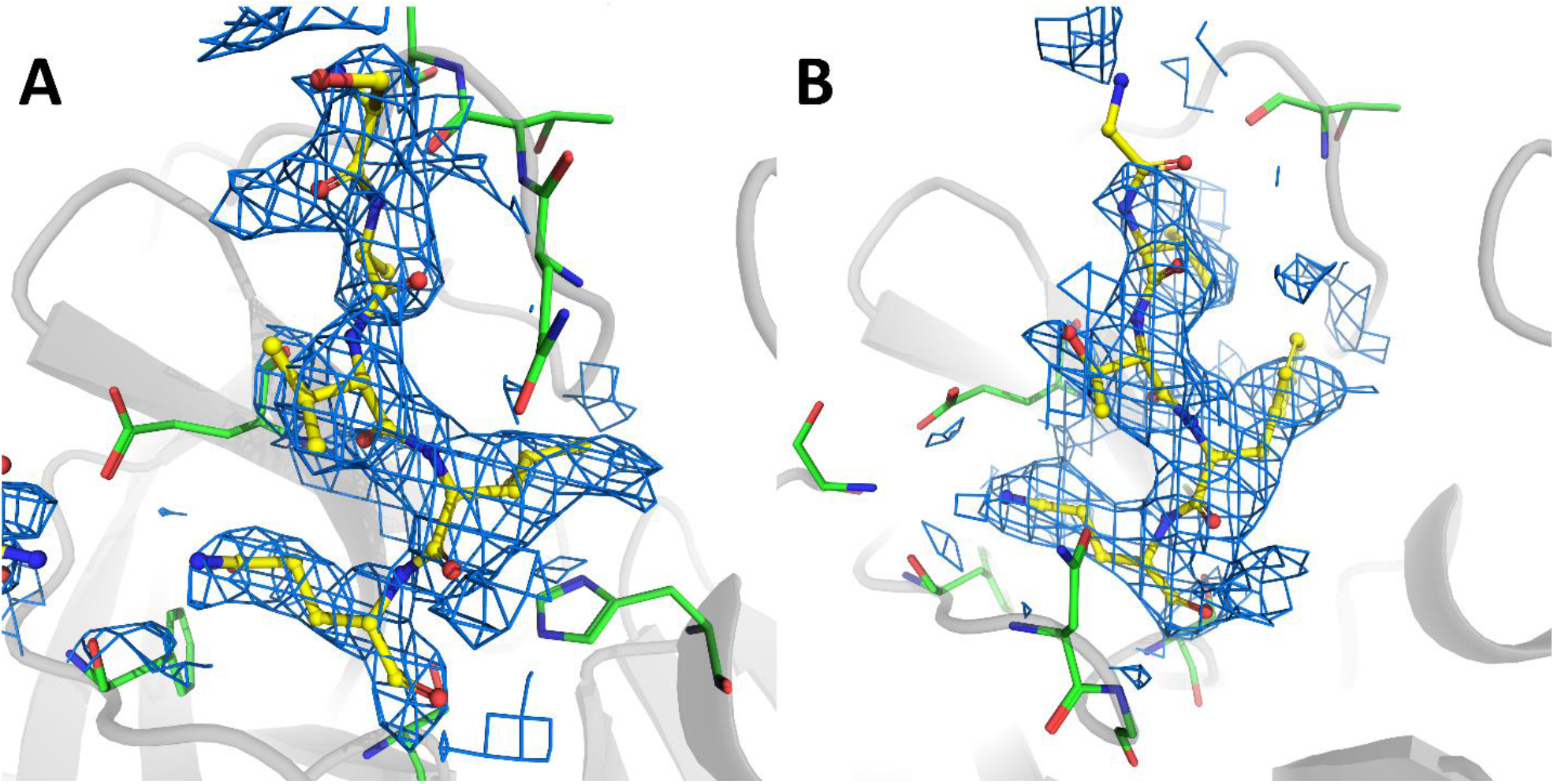
(A) Simulated annealing OMIT maps of N-terminal residues bound to C145S M^pro^ chain A (B) Simulated annealing OMIT maps of C-terminal residues bound to C145S M^pro^ chain B. OMIT maps were generated with RESOLVE (*42*). OMIT map density is contoured at 1.0 σ.

**Fig S8.**
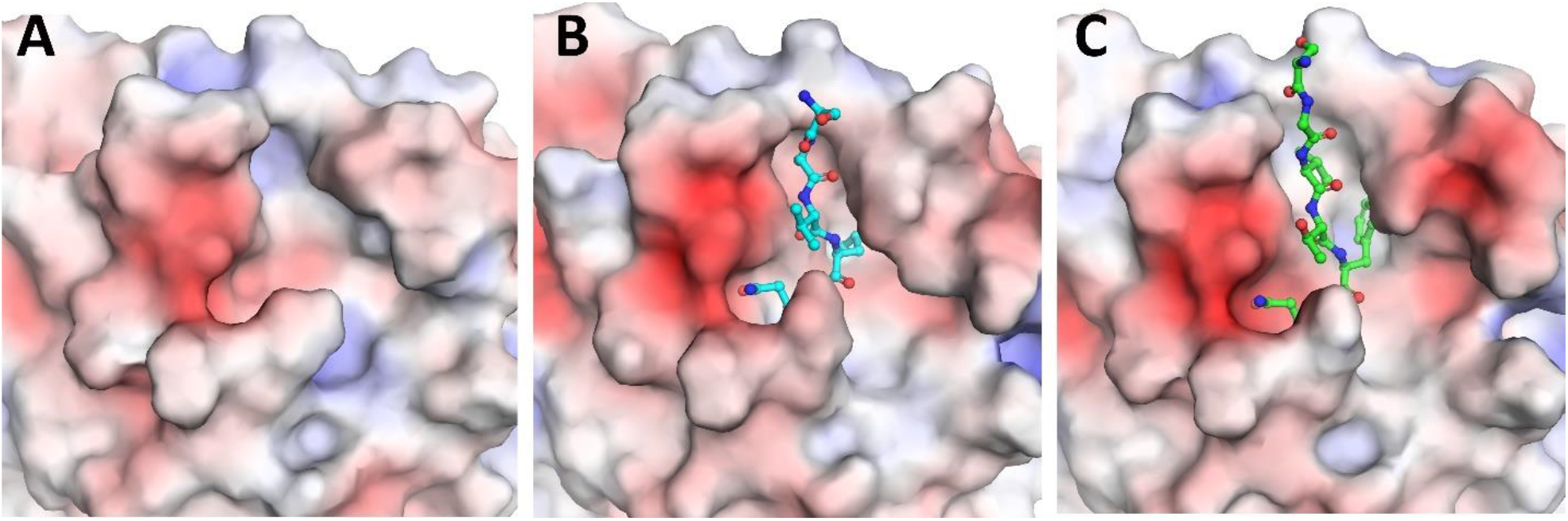
(a) Surface charge of M^pro^ active site. (b) Surface charge of C145A M^pro^ active site from chain A. N-terminal endogenous substrate is depicted as blue sticks. (c) Surface charge of C145A M^pro^ active site from chain B. C-terminal endogenous substrate is depicted as blue sticks. Surface charge was calculated with APBS (*43*).

**Fig S9.**
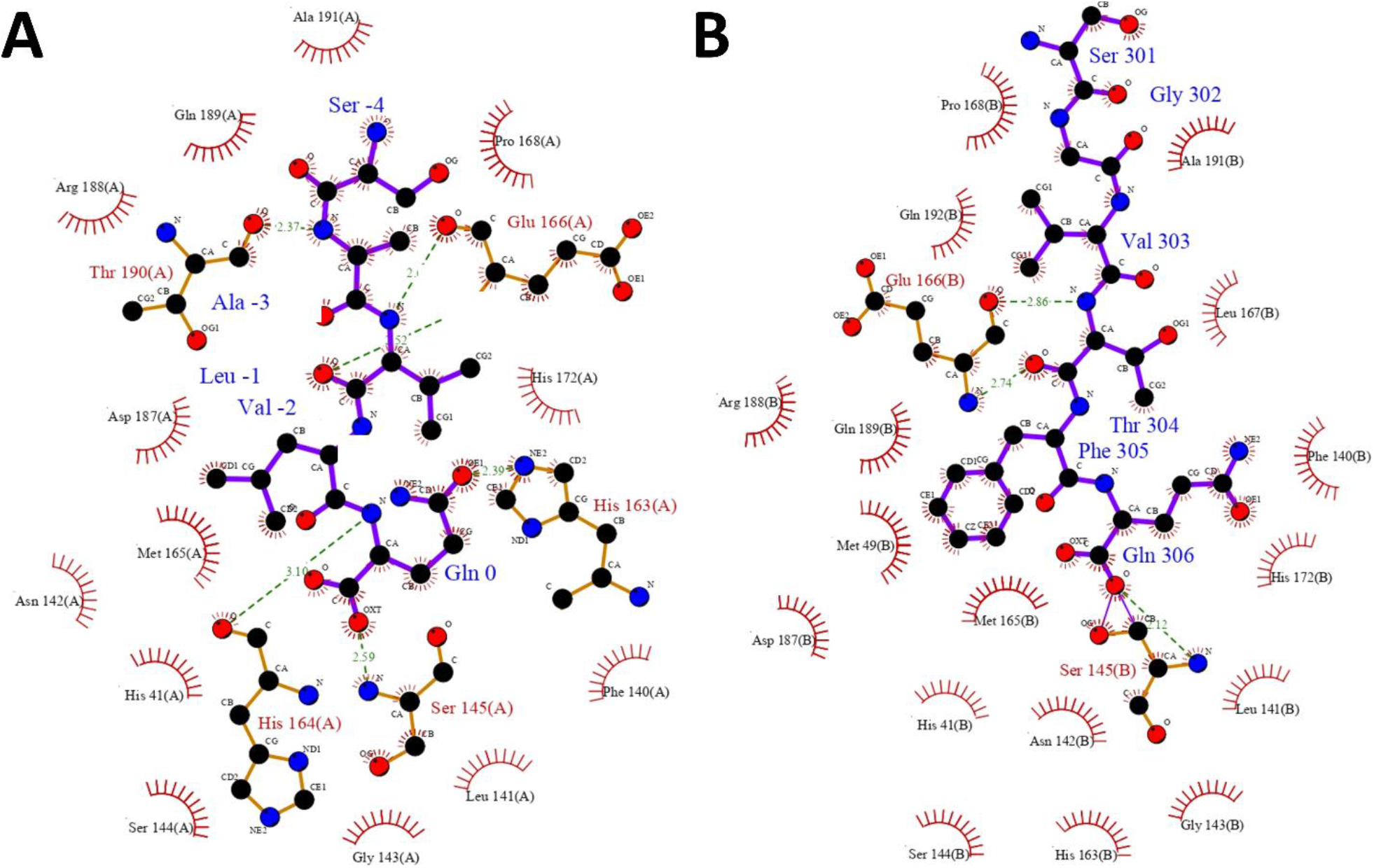
(a) Schematic drawing of interacting residues between C145A M^pro^ and N-terminal endogenous substrate. (b) Schematic drawing of interacting residues between C145A M^pro^ and C-terminal endogenous substrate. Figure was generated with LigPlot (*44*).

**Fig S10.**
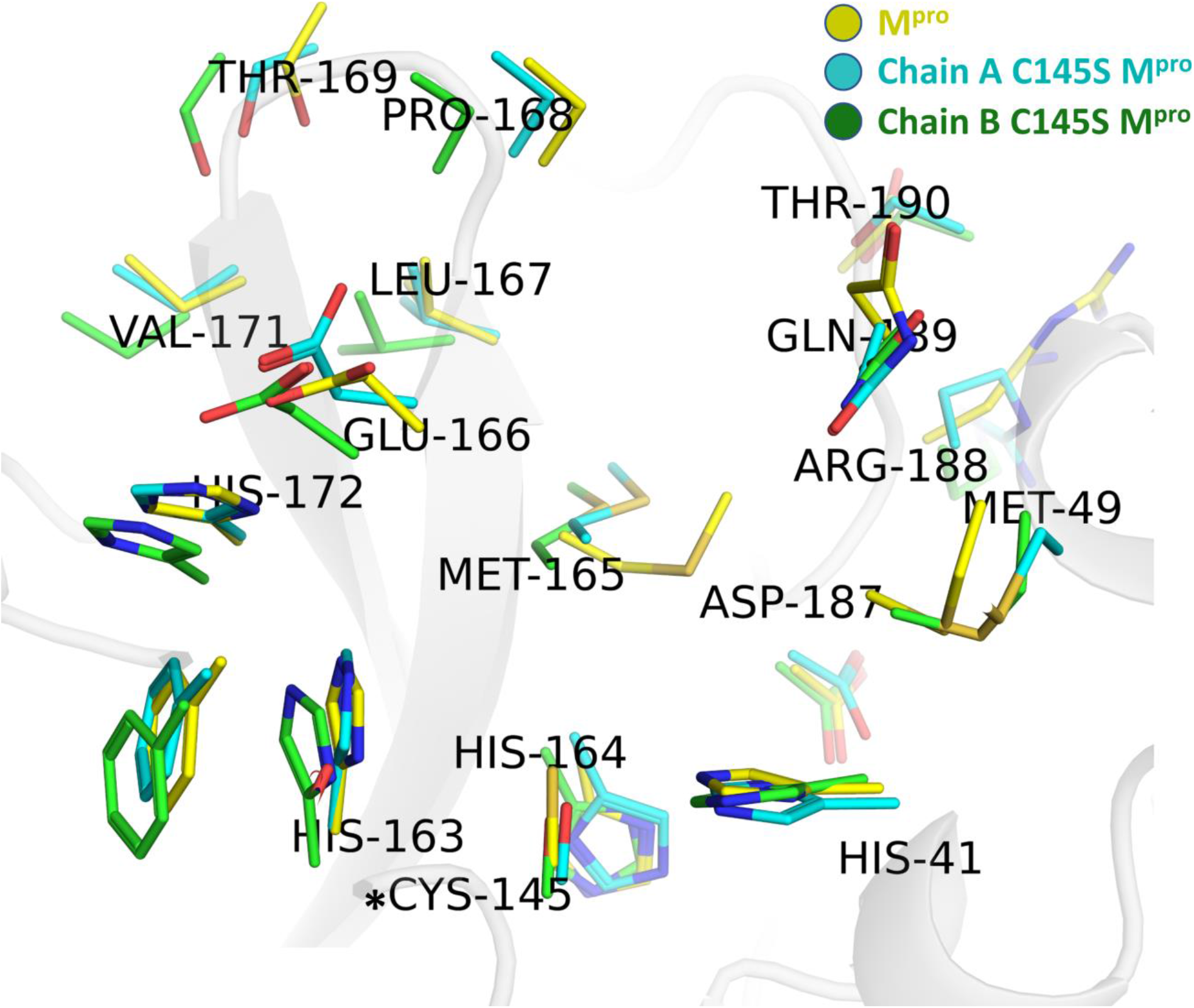
Superposition of the active sites from M^pro^ (yellow), C145A M^pro^ chain A (cyan) and C145A M^pro^ chain B (green). *CYS145 is SER145 for C145A M^pro^ chains A and B.

**Fig S11.**
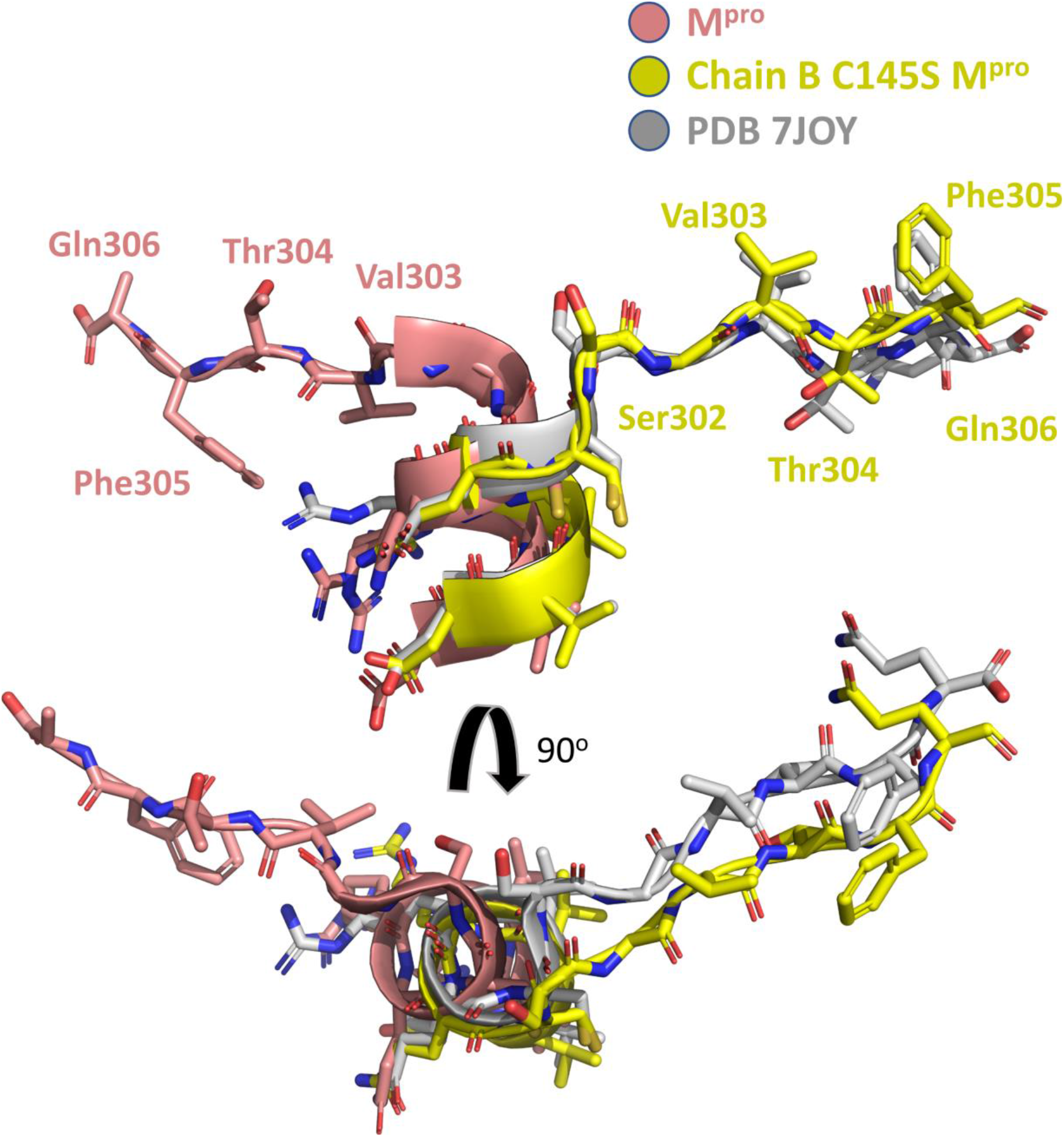
Superposition of C-terminal residues from M^pro^ (salmon), C145A M^pro^ chain B (yellow) and PDB 7JOY chain B (grey).

**Fig S12.**
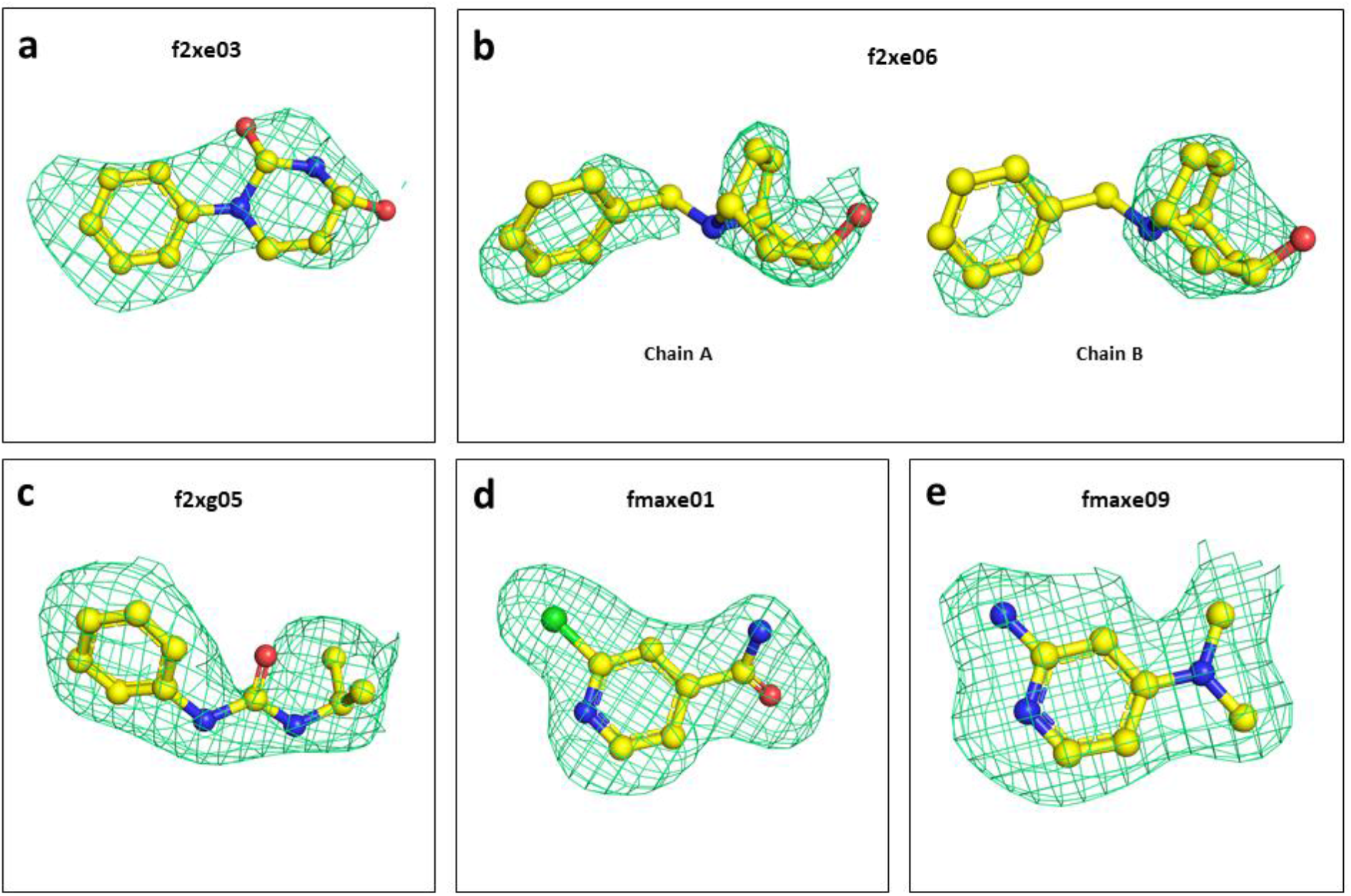
Polder maps of identified fragments for IMT M^pro^, showed with σ = 1.0.(*45*)

**Scheme S1.**
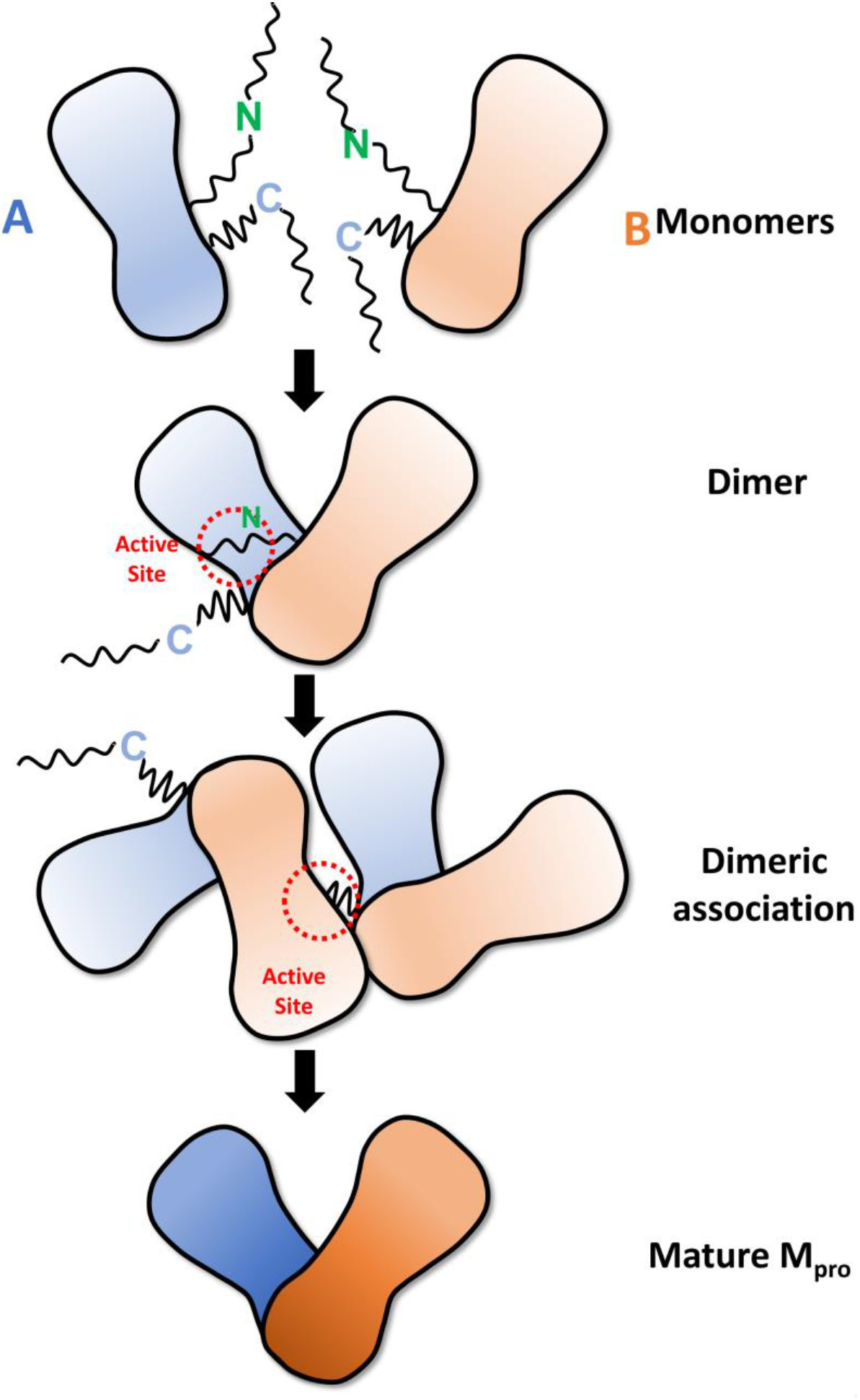
Schematic showing steps of SARS-CoV-2 M^pro^ maturation process. The immature monomeric form of M^pro^ form an intermediary dimer during cis-cleavage of N-terminal. With a mature N-terminal, the M^pro^ is then trans-cleaved by the dimeric association with another dimer, producing full mature M^pro^. M^pro^chain A and B are shown in blue and salmon, respectively. N and C terminals are marked in green and blue, respectively. Polyprotein is shown as black lines.

**Table S1.** Processing parameters and statistics of datasets 01, 02 and 03, used for beamline testing and structure determination of IMT M^pro^.

|  | **data01** | **data02** | **data03** |
| --- | --- | --- | --- |
| **Resolution** | 72.24 -1.87 (1.97-1.87) | 56.35-1.84 (1.94-1.84) | 67.60-1.83 (1.93-1.83) |
| **Space groups** | *P*2<sub>1</sub>2<sub>1</sub>2<sub>1</sub> | *P*2<sub>1</sub>2<sub>1</sub>2<sub>1</sub> | *P*2<sub>1</sub>2<sub>1</sub>2<sub>1</sub> |
| **Unit Cells (Å)** | 67.6, 102.0,102.3 | 67.5, 101.9, 102.1 | 67.6, 101.5, 102.25 |
| ***R*<sub>merge</sub>** | 0.054 (0.574) | 0.053 (0.634) | 0.059 (0.661) |
| ***R*<sub>meas</sub>** | 0.064 (0.683) | 0.063 (0.759) | 0.070 (0.790) |
| ***R*<sub>pim</sub>** | 0.034 (0.365) | 0.034 (0.410) | 0.037 (0.428) |
| ***R*<sub>merge</sub> in top intensity bin** | 0.033 | 0.032 | 0.035 |
| **Total number unique** | 57163 (8596) | 56445 (6967) | 58535 (6727) |
| ***I*/σ*I*** | 16.7 (2.7) | 16.3 (2.4) | 16.1 (2.4) |
| **CC<sub>(1/2)</sub>** | 0.999 (0.900) | 0.999 (0.872) | 0.999 (0.852) |
| **Completeness** | 95.7 (99.9) | 91.4 (78.5) | 94.0 (75.1) |
| **Multiplicity** | 6.5 (6.4) | 6.4 (6.2) | 6.7 (6.4) |

**Table S2.** Data collection and refinement statistics.

|  | M<sup>pro</sup> | IMT M<sup>pro</sup> | C145S M<sup>pro</sup> |
| --- | --- | --- | --- |
| *Data collection* |  |  |  |
| Space group | C2<sub>1</sub> | P2<sub>1</sub> 2<sub>1</sub> 2<sub>1</sub> | P2<sub>1</sub> |
| Cell dimensions |  |  |  |
| *a, b, c* (Å) | 113.0, 52.8, 44.7 | 67.6, 101.8, 102.2 | 58.6, 79.3, 62.8 |
| $\alpha, \beta, \gamma$ (°) | 90, 102.8, 90 | 90, 90, 90 | 90, 106.5, 90 |
| Resolution range (Å) | 55.1-1.46 (1.51-1.46) | 29.1-1.6 (1.65-1.6) | 60.2-2.8 (2.9-2.8) |
| Unique Reflections | 42249 (5935) | 93647 (4597) | 13665 (1976) |
| Multiplicity | 3.4 (3.3) | 18.4 (15.8) | 3.7 (3.3) |
| Completeness (%) | 94.9 (91.2) | 99.9 (99.7) | 99.36 (97.9) |
| *I*/σ*I* | 12.8 (2.4) | 8.8 (0.7) | 3.3 (1.1) |
| *R*<sub>p.i.m.</sub> (%)<sup>1</sup> | 0.031 (0.298) | 0.023 (0.90) | 0.15 (0.83) |
| CC<sub>1/2</sub><sup>2</sup> | 1.0 (0.46) | 1.0 (0.49) | 0.96 (0.38) |
| *Refinement* |  |  |  |
| *R*<sub>work</sub>/*R*<sub>free</sub><sup>3</sup> | 0.16/0.18 | 0.21/0.23 | 0.20/0.24 |
| Number of atoms |  |  |  |
| waters | 328 | 454 | - |
| Protein residues | 306 | 612 | 617 |
| RMS(bonds) (Å) | 0.008 | 0.014 | 0.011 |
| RMS(angles) (°) | 1.17 | 1.68 | 1.39 |
| Ramachandran favored (%) | 98.68 | 96.84 | 96.56 |
| Ramachandran outliers (%) | 0.33 | 0.17 | 0.33 |
| Clashscore<sup>4</sup> | 4.6 | 1.59 | 12.22 |
| Average *B*-factors (Å<sup>2</sup>) |  |  |  |
| Macromolecules | 24.7 | 37.28 | 48.22 |
| Solvent | 34.9 | 50.06 | - |
| PDB code | 7KPH | 7KFI | 7KVG |

<sup>1</sup> $R_{p.i.m.} = \sum_{hkl} \{1/[N(hkl) - 1]\}^{1/2} \times \sum_j |I_i(hkl) - \langle I(hkl) \rangle| / \sum_{hkl} \sum_j I_i(hkl)$ (46).
<sup>2</sup> CC<sub>1/2</sub> is the correlation coefficient determined by two random half data sets (47)
<sup>3</sup> $R_{work} = \sum_{hkl} |F_o(hkl) - F_c(hkl)| / \sum_{hkl} F_o(hkl)$ . *R*<sub>free</sub> was calculated for a test set of reflections (5%) omitted from the refinement.
<sup>4</sup> Clashscore is the number of clashes calculated for the model per 1000 atoms (48).

**Table S3.** Data collection and refinement statistics for observed fragments. Statistics for the highest-resolution shell are shown in parentheses.

|  | **f2xe03** | **f2xe06\*** | **f2xg05** | **fmaxe01** | **fmaxe09** |
| --- | --- | --- | --- | --- | --- |
| **Resolution range**<br>(Å) | 51.3 - 2.8 (2.9 - 2.8) | 56.1 - 2.2 (2.3 - 2.2) | 37.6 - 2.6 (2.7 - 2.6) | 72.09 - 2.09 (2.1 - 2.09) | 72.02 - 2.12 (2.19 - 2.12) |
| **Space group** | *P2<sub>1</sub>2<sub>1</sub>2<sub>1</sub>* | *P2<sub>1</sub>2<sub>1</sub>2<sub>1</sub>* | *P2<sub>1</sub>2<sub>1</sub>2<sub>1</sub>* | *P2<sub>1</sub>2<sub>1</sub>2<sub>1</sub>* | *P2<sub>1</sub>2<sub>1</sub>2<sub>1</sub>* |
| **Unit cell (a,b,c;Å)** | 67.7, 102, 102.6 | 67.3, 101.6, 101.9 | 67.5, 101.0, 102.1 | 67.6, 101.9, 101.9 | 67.6, 101.4, 102.2 |
| **Unique reflections** | 96410 (11291) | 43094 (499) | 40900 (4004) | 41113 (4161) | 39479 (3970) |
| **Completeness (%)** | 93.60 (99.60) | 61.77 (7.23) | 99.84 (100.0) | 96.77 (99.93) | 97.19 (99.87) |
| **R-work** | 0.2191 | 0.2410 | 0.2141 | 0.1923 | 0.2004 |
| **R-free** | 0.2605 | 0.2700 | 0.2658 | 0.2263 | 0.2244 |
| **Number of non-hydrogen atoms** | 4783 | 4872 | 4992 | 5017 | 4817 |
| **macromolecules** | 4675 | 4663 | 4681 | 4680 | 4643 |
| **ligands** | 66 | 77 | 66 | 119 | 69 |
| **solvent** | 42 | 132 | 245 | 218 | 105 |
| **Protein residues** | 604 | 603 | 605 | 604 | 600 |
| **RMS(bonds)** | 0.007 | 0.007 | 0.007 | 0.009 | 0.014 |
| **RMS(angles)** | 1.28 | 0.87 | 1.10 | 1.35 | 1.81 |
| **Ramachandran favored (%)** | 97.00 | 95.49 | 96.84 | 96.83 | 97.99 |
| **Ramachandran allowed (%)** | 3.00 | 4.34 | 3.16 | 2.83 | 1.85 |
| **Ramachandran outliers (%)** | 0.00 | 0.17 | 0.00 | 0.33 | 0.17 |
| **Rotamer outliers (%)** | 0.00 | 1.35 | 0.58 | 0.38 | 1.94 |
| **Clashscore** | 8.31 | 6.94 | 7.02 | 6.40 | 8.79 |
| **Average B-factor** | 50.42 | 29.16 | 52.56 | 42.83 | 51.25 |
| **macromolecules** | 50.15 | 29.04 | 52.31 | 42.17 | 51.01 |
| **ligands** | 74.64 | 45.08 | 71.73 | 63.79 | 72.19 |
| **solvent** | 41.66 | 23.86 | 52.16 | 45.55 | 47.99 |
| **PDB CODE** | 7LFE | 7LDX | 7LFP | 7KVL | 7KVR |

\*processed with STARANISO (39)

**Table S4:**
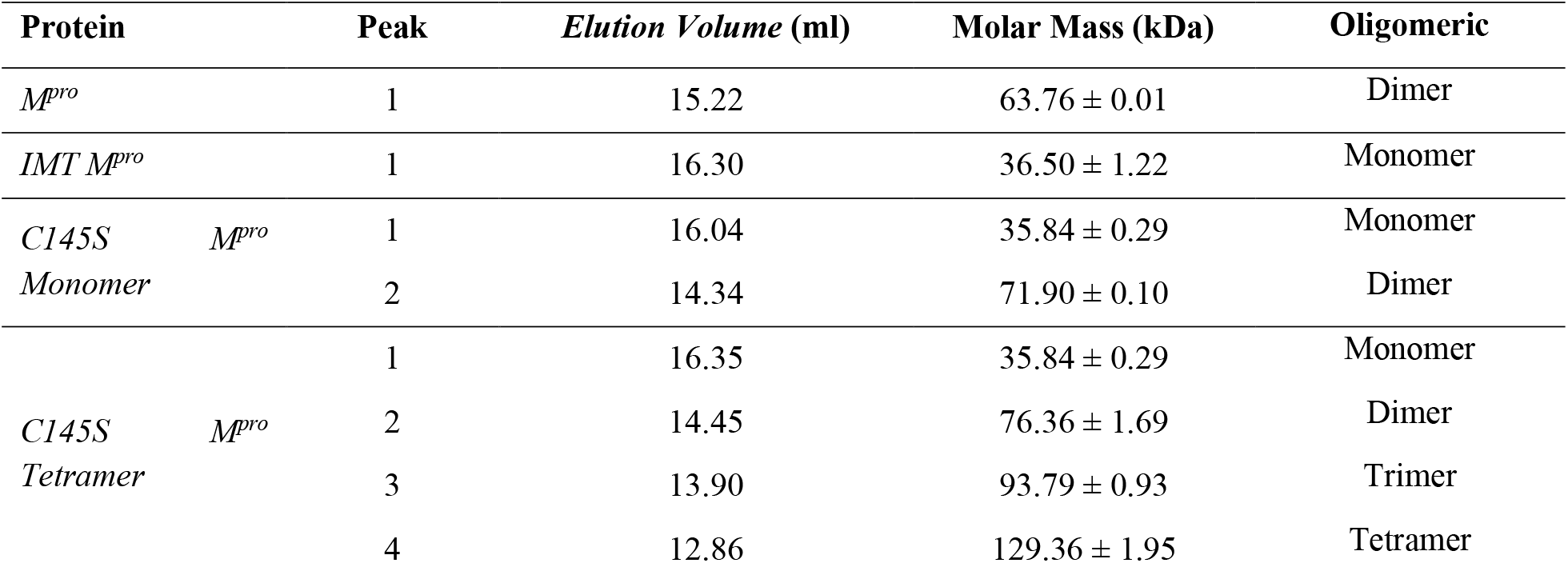
Molecular Mass and Elution Volume for the each observed peak in the SEC-MALS profiles of SARS-CoV-2 M^pro^ constructs.

**Table S5.**
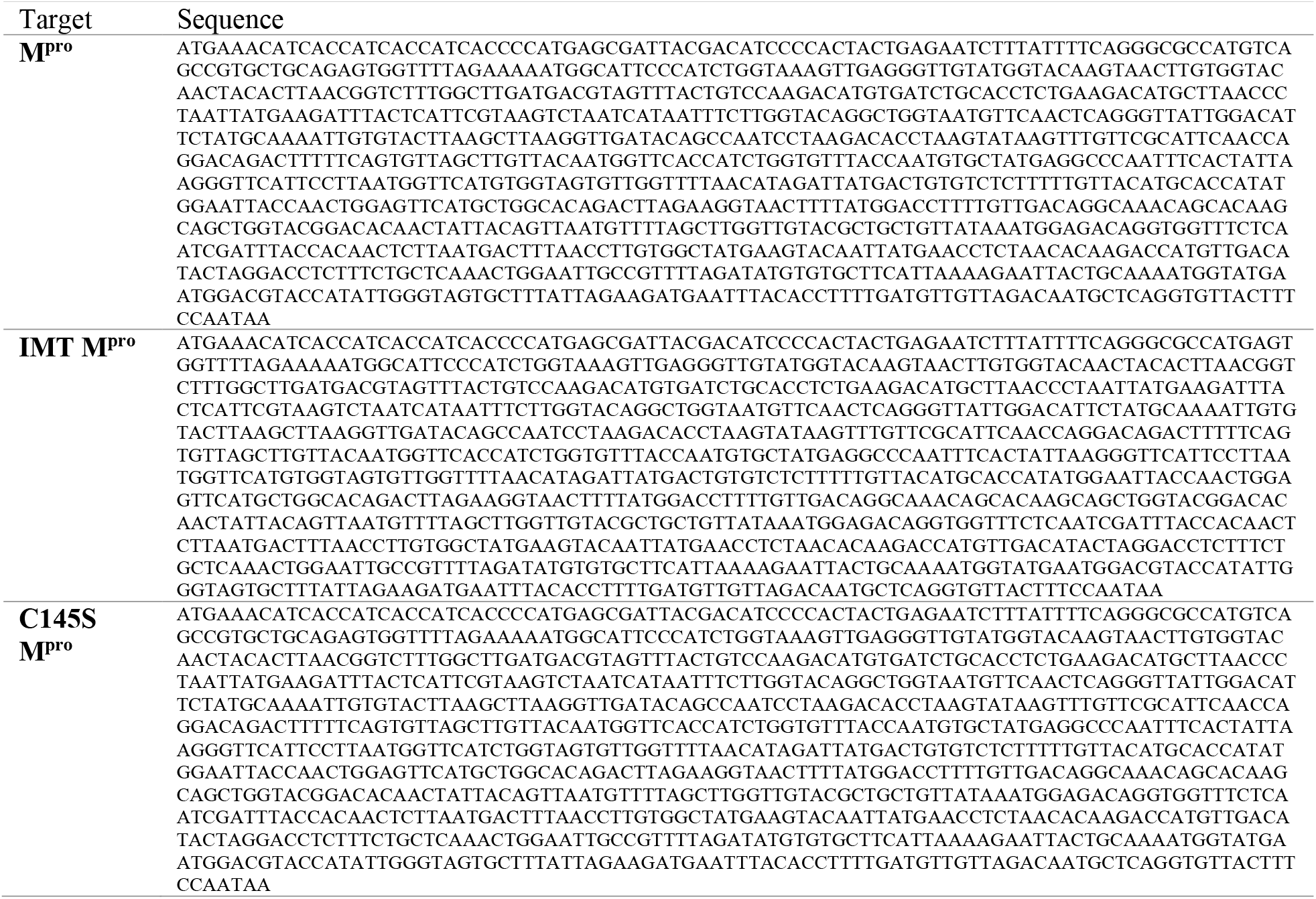
Sequences expressed of each M^pro^ construct

